# A Family of Glycosylated Macrolides Selectively Target Energetic Vulnerabilities in Leukemia

**DOI:** 10.1101/2021.05.31.445492

**Authors:** Benjamin J. Reisman, Hui Guo, Haley E. Ramsey, Madison T. Wright, Bradley I. Reinfeld, P. Brent Ferrell, Gary A. Sulikowski, W. Kimryn Rathmell, Michael R. Savona, Lars Plate, John L. Rubinstein, Brian O. Bachmann

## Abstract

Cancer cells have long been recognized to exhibit unique bioenergetic requirements. The apoptolidin family of glycomacrolides are distinguished by their selective cytotoxicity towards oncogene transformed cells, yet their molecular mechanism remains uncertain. We used photoaffinity analogs of the apoptolidins to identify the F1 subcomplex of mitochondrial ATP synthase as the target of apoptolidin A. CryoEM of apoptolidin and ammocidin-ATP synthase complexes revealed a novel shared mode of inhibition that was confirmed by deep mutational scanning of the binding interface to reveal resistance mutations which were confirmed using CRISPR-Cas9. Ammocidin A was found to suppress leukemia progression *in vivo* at doses that were tolerated with minimal toxicity. The combination of cellular, structural, mutagenesis, and *in vivo* evidence define the mechanism of action of apoptolidin family glycomacrolides and establish a path to address OXPHOS-dependent cancers.

---

The development of effective cancer therapeutics is constrained by the need to identify targets and drugs that selectively eliminate cancer cells with limited toxicity to healthy cells. One approach to addressing this challenge is to identify functional dependencies that are unique to cancer; even an ‘essential’ targets is present in non-cancerous cells, the unique demands of cancer cells create vulnerabilities that can be leveraged therapeutically^1^. Alterations in bioenergetics have long been recognized as a hallmark of cancer^2–4^, with Warburg’s observation that some cancer cells convert glucose to lactate in the presence of adequate oxygen – aerobic metabolism. Although Warburg metabolism predominates in many cancers, there is increasing recognition that some cancers and subsets of cancer cells are dependent on mitochondrial oxidative phosphorylation (OXPHOS) to serve both bioenergetic and biosynthetic needs^5–7^. Notably, OXPHOS dependence appears to be a recurrent vulnerability of cancer stem cells (CSCs) and leukemia stem cells (LSCs), which are often responsible for treatment failure^8–11^. Despite these vulnerabilities, outside of a small group of genetically defined tumor subsets^12^, therapeutics designed for metabolic targets in cancer have largely fallen short of their great promise^13^.

The apoptolidin family of glycosylated macrolides (glycomacrolides) was discovered in a series of phenotypic screens for compounds that induce apoptosis selectively in oncogene-transformed cells. Apoptolidin A, produced by *Nocardiopsis* sp., was discovered to selectively induce apoptosis in E1A transformed rat fibroblasts^14,15^, while ammocidin A, a structurally related glycomacrolide produced by *Saccharothrix* sp., was discovered in a screen for compounds that selectively induce apoptosis in HRas-G12V transformed BaF3 cells^16,17^. When evaluated in the NCI-60 screening program, apoptolidin A was noted to be among the top 0.1% most cell line selective cytotoxic agents tested at that time, and early studies of its mechanism of action suggested that it targets ATP synthase, a mitochondrial membrane protein complex involved in oxidative phosphorylation ^18^. These findings prompted multiple groups to develop new analogs via fermentation^19–24^, semi-synthesis^25,26^, and total synthesis^27–30^. Structure-activity relationship (SAR) studies of these analogs supported a role for OXPHOS inhibition at ATP synthase in the mechanism of apoptolidin family compounds, but subtle differences in the bioactivity of apoptolidins compared to other classical ATP synthase inhibitors such as oligomycin suggested a distinct biochemical target^26,31^. Given the remarkable selectivity of apoptolidin family compounds *ex vivo* and emerging evidence of the importance of OXPHOS in cancer, identification of the apoptolidin target could reveal new targetable dependencies in cancer metabolism. In this study, we applied photoaffinity probes of apoptolidin family macrolides and electron cryomicroscopy (cryoEM) to identify their specific target and binding site on the F_1_, rather than F_O_, subcomplex of ATP synthase. Resistance mutations uncovered by deep mutational scanning eliminated both binding and cytotoxicity, confirming the mechanistic role for ATP synthase in their selective cytotoxicity. Ammocidin A was able to suppress leukemia development in mouse xenografts demonstrating that ATP synthase can be effectively targeted in OXPHOS dependent cancers.

## Results

### Identification of the F_1_ subcomplex of ATP synthase as the target of glycomacrolides

The apoptolidin family of glycomacrolides, including apoptolidin A (**1**), apoptolidin H (**3**), and ammocidin A (**5**) all incorporate a 20-membered macrolide conjugated to three deoxy sugar units and are produced by taxonomically related actinobacteria. While superficially structurally similar, there are multiple differences in the functionality of the macrolide cores, decorating sugars, and in the positioning of glycosyl units. In order to identify the target of this family, we synthesized photoaffinity probe analogs of apoptolidin family macrolides (Fig. 1a). The C9 2-deoxyglucose was selected as the site of modification as previous studies had demonstrated that it could tolerate modification with retention of nanomolar cytotoxicity^32^. Apoptolidin A PA (**2**) incorporates a diazirine, which provides for *in situ* photoreactive crosslinking when exposed to UV-A light^33^, as well as an alkyne, which allows for post-treatment ligation of fluorophores or affinity enrichment tags using click chemistry^34^. The MV-4-11 acute myeloid leukemia cell line was selected as a primary model system for target identification based on its high sensitivity to apoptolidin family compounds The activity of (**2**) was comparable to (**1**) with regards to both cytotoxicity (Extended Data Fig. 1a) and suppression of S6 [Ser235/236] phosphorylation (pS6 – Extended Data Fig. 1b), a marker of activation of the AMPK pathway which has been implicated in the mechanism of action of apoptolidin^31^. When applied to adherent, apoptolidin sensitive, H292 cells and probed with rhodamine azide after fixation, (**2**) exhibited the expected mitochondrial localization (Extended Data Fig. 1d) consistent with previous studies using fluorescent live cell apoptolidin probes^32^.

**Fig. 1.**
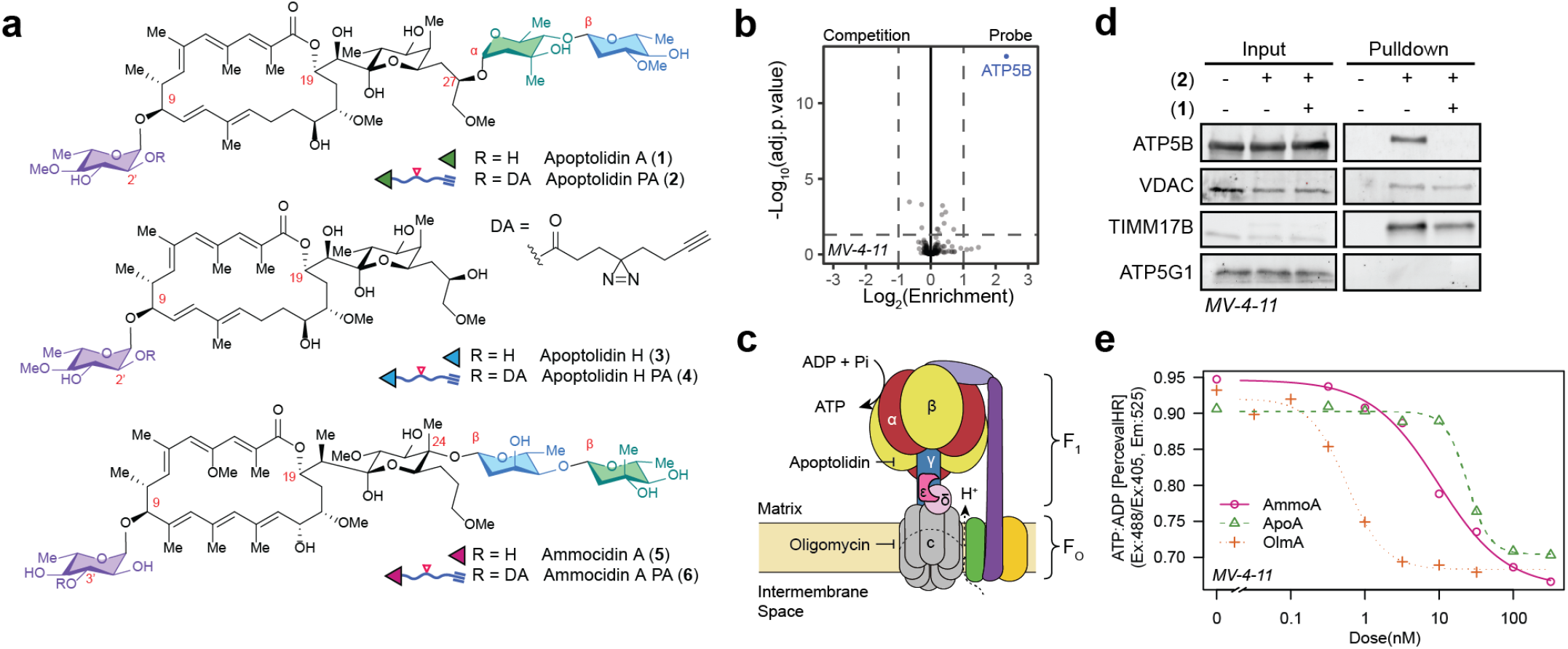
Identification of F_1_ subcomplex of ATP synthase as the target of apoptolidin A. **a,** Structures of natural products and probes used in this study with key differences between apoptolidin and ammocidin highlighted. **b**, Volcano plot of identified proteins from TMT multiplexed quantitative proteomics of (**2**) (probe) vs (**1**) + (**2**) (competition) treated MV-4-11 cells in duplicate, adjusted p-values calculated using Limma. **c,**Illustration of the mitochondrial ATP synthase; **d,** Immunoblot of MV-4-11 cells treated with (**1**) [1 μM] and/or (**2**) [20 μM] showing specific enrichment of ATP5B, non-specific enrichment of VDAC and TIMM17B, and no enrichment of ATP5G1. **e,** Inhibition of ATP synthesis after 16 h treatment with AmmoA, ApoA, or OlmA measured in MV-4-11 leukemia cells expressing the ATP:ADP reporter Perceval HR.

To identify apoptolidin family binding proteins, MV-4-11 cells were treated with DMSO, probe (**2**) alone, or (**2**) combined with various potential competitors (Extended Data Fig. 1e). Proteins were photocrosslinked prior to lysis and then conjugated via azide-alkyne *Huisgen* cycloaddition to a trifunctional TAMRA-azide-desthiobiotin reagent, which facilitated both gel-based fluorescent profiling and subsequent affinity enrichment. Gel-based profiling of (**2**) adducts (Extended Data Fig. 1f) revealed concentration-dependent labelling of a ~50 kDa target with an apparent EC_50_ of ~10 nM. Adduction of the 50 kDa target was eliminated by competition with excess apoptolidin A (**1**) as well as ammocidin A (**3**) suggestive of a shared putative binding site for the two glycomacrolides. Apoptolidin H (**3**), lacking the C-27 disaccharide and known to have reduced cytotoxicity relative to apoptolidin A^22^, exhibited decreased competitive displacement of apoptolidin A PA (**2**). Furthermore, oligomycin did not exhibit any competition, suggesting that the 50 kDa target engaged by apoptolidins and ammocidin A was not the target of oligomycin, as previously proposed^35^. To confirm similar adduction specificity across the family we synthesized photoaffinity probes apoptolidin H PA (**4**) and ammocidin A PA (**6**), which also retain cytotoxicity similar to the parent compounds (Extended Data Fig. 1c). Gel-based profiling of adducted proteins demonstrated the same 50 kDa band (Extended Data Fig. 1g,h) Similar binding patterns were observed in additional cell lines (Extended Data Fig. 1i-k) including HRas-G12V transformed BaF3s and H292 lung cancer cells, similar to those used in previous studies on apoptolidin family glycomacrolides^16,25^. The trienoate and C21 hemiketal functional groups of apoptolidin A are potential electrophiles that could allow apoptolidin to act as a covalent inhibitor. However, irreversible binding to the 50 kDa target was dependent on both the diazirine and exposure to UV light, suggesting that covalent modification of the target does not play a role in target engagement in naturally occurring apoptolidin family macrolides (Extended Data Fig. 3b).

To identify the putative targets of apoptolidin A in the MV-4-11 proteome, affinity enrichment was performed after *in situ* photocrosslinking of (**2**) with or without excess apoptolidin A (**1**). Subsequent to lysis and cycloaddition to TAMRA-azide-desthiobiotin, samples were affinity enriched using streptavidin resin and labeled for multiplexed mass spectrometry quantification using isobaric tandem mass tags (TMT) (Extended Data Fig. 2b). Quantitative proteomics on the enriched tagged samples revealed a single, highly enriched target relative to the competition control: ATP5B, the β subunit of ATP synthase (Fig. 1b). ATP synthase is a ~600 kDa multiprotein complex responsible for converting the electrochemical proton gradient generated by the electron transport chain into chemical energy by the synthesis of ATP from ADP and phosphate. ATP synthase is composed of two subcomplexes: the membrane-embedded F_O_ region, which contains the proton channel and is the target of oligomycin, and the soluble F_1_ region, which contains the catalytic sites (Fig. 1c). The F_1_ subcomplex is composed of five distinct subunits with the stoichiometry α_3_β_3_γδε and is largely conserved across the kingdoms of life^36^. The three catalytic sites are located at the interfaces of the α and β subunits with the catalytic residues localized to the β subunit, which is encoded by the gene *ATP5F1B* in humans. The three catalytic αβ sites are found in different conformations designated by their nucleotide content as α_DP_β_DP_ (ADP-bound), α_TP_β_TP_ (ATP-bound), α_E_β_E_ (empty). During ATP synthesis, proton translocation through the F_O_ region induces rotation of the central rotor complex that drives ATP synthesis in the F_1_ region^37^. The enzyme can also catalyze ATP hydrolysis (ATPase), which causes the rotor to reverse direction and pump protons from the matrix to the intermembrane space.

Several non-specific adducts were enriched in the competition conditions compared to vehicle (Extended Data Fig. 2c,d). These include the mitochondrial membrane proteins VDAC and TIMM17B, as well as other membrane proteins, many of which are documented non-specific targets of alkyl diazirines^38^. Immunoblotting confirmed the specific interaction of (**2**) with the β subunit of ATP synthase and nonspecific interactions with VDAC, and no interaction with ATP5G1, the c subunit in the FO region that is target of oligomycin (Fig. 1d). Several attempts were made to identify the adduction site of (**2**) on the β subunit using shotgun proteomics, but these were unsuccessful, likely due to the large size of the theoretical adduct. β subunit binding by (**2**) was also confirmed by direct in-gel digestion of Coomassie dye stained bands after affinity enrichment (Extended Data Fig. 2e). Finally, binding specificity to the β subunit was also confirmed in HEK-293 landing pad (HEK293-LP) cells expressing transgenic *ATP5F1B* with a C-terminal FLAG-tag (Extended Data Fig. 1l).

ATP synthase had been proposed as a target of apoptolidin family glycomacrolides based on similar cytotoxicity profiles to oligomycin in the NCI-60 screening program^18^. Due to their common general classification as macrolides, apoptolidin and oligomycin were proposed to share a SAR model and both bind to the F_O_ region of ATP synthase^39^. More recent studies questioned whether the ATP synthase FO region is truly the mechanistic target of apoptolidin due to the discrepancy between *in vitro* (low μM) and *ex vivo* (low nM) activity^26^ and different effects on cell signaling^31^. These data demonstrate that apoptolidin and oligomycin act via distinct mechanisms at different binding sites on ATP synthase. Of note, apoptolidin and ammocidin represent the first known bacterial compounds to target the F_1_ ATP synthase, as all previously described F_1_ inhibitors are of fungal origin^40^, while bacterial ATP synthase inhibitors such as oligomycin target the F_O_ region^41^.

To evaluate whether apoptolidin was acting as an inhibitor of ATP synthase at the cytotoxic concentration, ATP synthesis was evaluated in MV-4-11 cells expressing the intracellular ADP/ATP reporter, Perceval HR^42^. Treatment with apoptolidin A, ammocidin A, or oligomycin A elicited an increase in ADP/ATP ratio at concentrations comparable with their cytotoxicity (Fig. 1e). *In vitro*, apoptolidin A and ammocidin A exhibited similar low nano-molar inhibition of ATPase activity by purified *Saccharomyces cerevisiae* ATP synthase (Extended Data Fig. 1m). Competition experiments with known F_1_ binding inhibitors revealed uncompetitive binding with aurovertin and a lack of binding to the efrapeptin-bound conformation (Extended Data Fig. 3c). Kinetics studies in isolated mouse liver mitochondria and in yeast F_1_ subcomplexes under non-saturating conditions revealed mixed inhibition (Extended Data Fig. 3e,f) consistent with an allosteric binding site distal to the catalytic sites at the interfaces of the α and β subunits.

Surprisingly, pre-treatment of cells with the uncoupling agents carbonyl cyanide *p*-trifluoromethoxyphenylhydrazone (FCCP) or 2,4-dinitrophenol (DNP) also eliminated adduction of (**2**) to the β subunit (Extended Data Fig. 3d). This observation suggested that either transport of (**2**) to the mitochondrial matrix was membrane potential dependent and/or the addition of uncouplers eliminated the apoptolidin binding site. Under uncoupled conditions, the decrease in mitochondrial matrix pH causes a conformational change in the inhibitory factor (ATPIF1, IF_1_) that allows it to bind to the F1 subcomplex and inhibit the ATPase activity of ATP synthase^43^. To test if IF_1_ binding was interfering with apoptolidin binding, CRISPR/Cas9 was used to knock out *ATPIF1* in K562 leukemia cells (Extended Data Fig. 3g). In this setting, FCCP eliminated binding of (**2**) to the β subunit in the parental line, but not in cells with *ATP5IF1* knocked out (Extended Data Fig. 3h), suggesting that IF_1_ prevents binding of apoptolidin to ATP synthase.

### Identification of the glycomacrolide binding site on ATP synthase via cryoEM

The novel mode of inhibition of glycomacrolides revealed by photoaffinity labeling and enzymatic assays prompted a search for their binding site by structure studies of *S. cerevisiae* ATP synthase. CryoEM and image analysis yielded 3D maps of the ATP synthase bound to ammocidin A and apoptolidin A at 3.1 and 3.4 Å resolutions, respectively, allowing construction of atomic models for the complexes. In order to investigate conformational changes induced in ATP synthase upon inhibitor binding, we also determined a structure of the yeast ATP synthase without either inhibitor under otherwise identical conditions, and refined the F_1_ region of the map to 4.2 Å resolution (Extended Data Fig. 4, Table S4).

Density corresponding to the inhibitor is found in the catalytic F1 region of ATP synthase (Fig. 2a, pink density). Despite the presence of three-fold pseudosymmetry in the F1 region, which provides three nearly-equivalent potential glycomacrolide binding sites, only a single inhibitor molecule was found to bind at the α_DP_β_DP_ interface. This binding site is formed mainly by subunits α_DP_, β_DP_, and γ, although neighboring β_TP_ and α_E_ subunits provide additional protein-inhibitor contacts (Fig. 2c). Despite the functional differences in the aglycone, attachment site, and sequence of glycosylation between apoptolidin A and ammocidin A, both molecules bind to ATP synthase in a strikingly similar manner (Fig. 2b, green density) and for clarity this discussion is focused on the higher-resolution ammocidin structure. The core macrolide is buried deeply inside the binding pocket and interacts with multiple hydrophobic residues of nearby subunits, while the head and tail sugar groups form fewer interactions with the protein (Fig. 2c, d). The structure indicates that introducing photoaffinity probes at the C-2’ position of the C-9 deoxysugar of apoptolidin, and the C-3’ position of ammocidin (Fig. 1a) is unlikely to interfere with drug binding, supporting the reliability of the photoaffinity labeling experiments described above. β_TP_-D386 is located close the C9 sugar and represents a likely site of photoaffinity adduct formation^38^. Together, these observations indicate that the high-affinity binding of ammocidin and apoptolidin to ATP synthase are achieved through extensive interactions between the core macrolides and their hydrophobic binding site in the enzyme (Fig. 2d). Comparison of the inhibitor-free and inhibitor-bound structures shows that inhibitor binding forces subunit β_DP_ to move outward to adopt a more “open” conformation (Fig. 2e, red arrows and Movie 1), while the conformation of the non-catalytic α_DP_ subunit remains largely unchanged except for the residues in direct contact with the inhibitor. This “open” conformation of β_DP_ is incompatible with nucleotide binding and explains why the only natural substrate found in the inhibitor-bound structures is a phosphate ion at the β_TP_ subunit (Fig. 2f). In addition to making β_DP_ incapable of binding substrates, the inhibitors likely also block the rotation of ATP synthase by interacting strongly with subunits α_DP_, β_DP_, and γ, preventing the conformational changes needed for ATP hydrolysis and thereby inhibiting the rotary catalytic mechanism of the enzyme^36^.

**Fig. 2.**
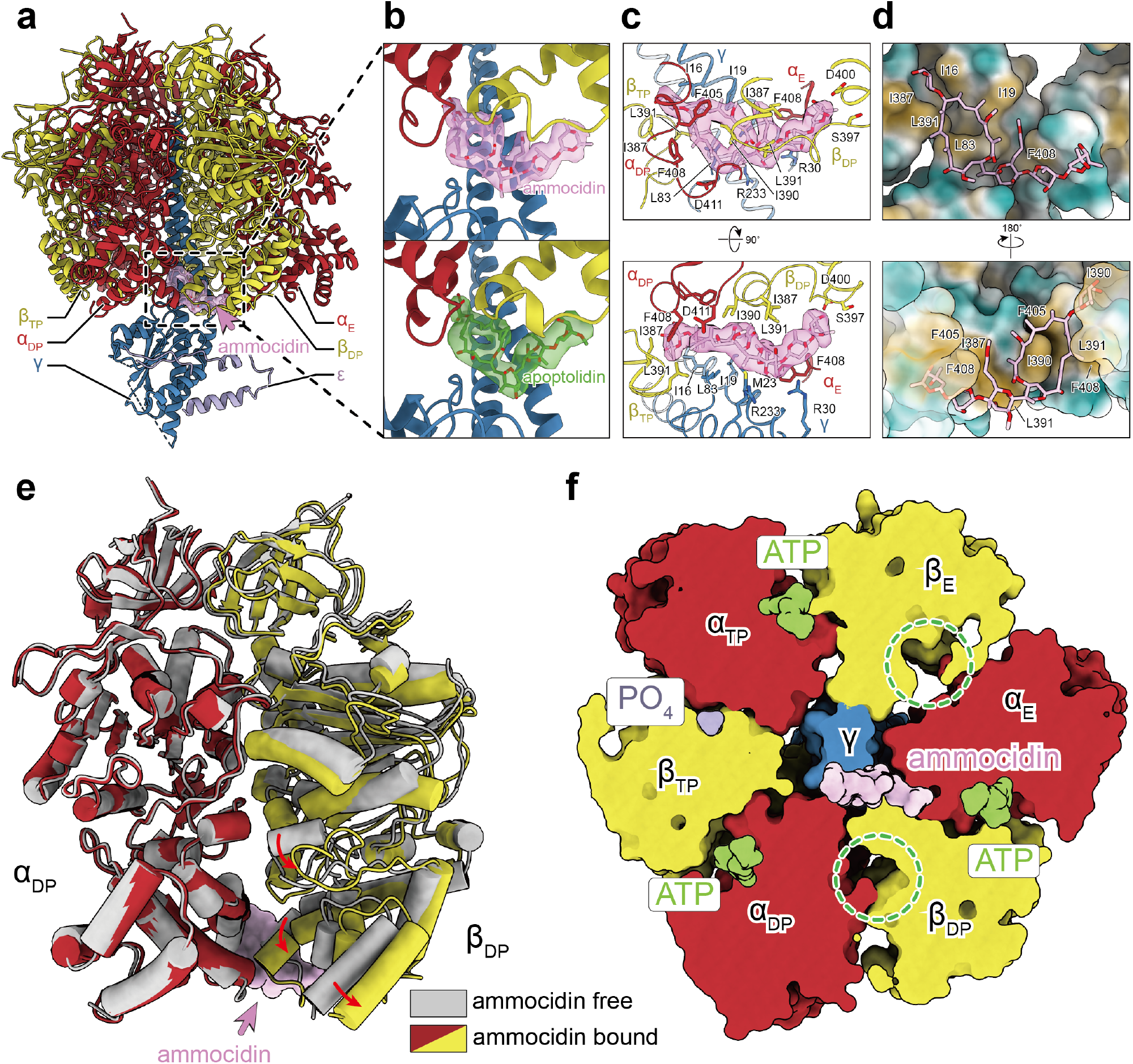
Structure of glycomacrolide bound yeast ATP synthase bound reveals a novel mode of inhibition. **a,** Atomic model of the F_1_ region of yeast ATP synthase bound to ammocidin, cryoEM density of ammocidin is shown in pink; **b,** Close-up view of the ammocidin (top) and apoptolidin (bottom) binding site; **c,** ammocidin binding site residues, **d,** Ammocidin binding pocket is largely formed with hydrophobic residues (yellow surface); **e,** Ammocidin binding induces conformational change of β_DP_ subunit (red arrows) and forces it to adopt a more “open” conformation; **f,** Cross section through the F_1_ region shows that β_DP_ does not bind to nucleotide (green circles) when bound to ammocidin.

### Deep mutational scanning of glycomacrolide binding site reveals mutations that confer resistance to apoptolidin and ammocidin

Having determined that the F_1_ region of ATP synthase is a target of apoptolidin A and ammocidin A, we next sought to determine whether binding to ATP synthase was essential for the cytotoxicity of apoptolidin by identifying one or more resistance mutations in the complex that prevent binding and cytotoxicity. Genomic analysis and multiple sequence alignment of ATP synthase genes from the apoptolidin producer *Nocardiopsis sp.* FU-40, revealed a substitution of phenylalanine at position ATP5B-L394 (Extended Data Fig. 5a) that we hypothesized would cause a clash that prevents binding. Introduction of ATP5B-L394F into MV-4-11 or K562 cell lines by CRISPR/Cas9 homology-directed repair (HDR)^44^ revealed decreased sensitivity to ammocidin A, but unexpectedly increased sensitivity to apoptolidin A compared to the parental line (Extended Data Fig. 5d,e). Comparing the binding poses of apoptolidin A to ammocidin A, ATP5B-L394 interacts most closely with the cyclic hemiketal that exhibits large differences in binding pose to account for the differing attachments of the disaccharide at C24 vs C27 (Extended Data Fig. 5f). Although this substitution does not provide cross resistance against both macrolides, the diverging effects on sensitivity supports the binding mode identified by cryoEM of the yeast enzyme.

In order to identify a mutation that confers complete resistance to both apoptolidin and ammocidin, a deep mutational scanning approach was taken, using the Bxb1 “landing pad” system^45^ to conduct a pooled screen of all possible missense mutations in the binding site. Mutagenesis targets were identified by selecting all residues within 4.5 Å of apoptolidin or ammocidin in the atomic models, identifying 21 residues across the α (5), β (8), and γ (8) subunits (Extended Data Fig. 6a), and a mutant library containing all 420 point mutants of the three genes was generated using nicking mutagenesis^46^ (Extended Data Fig. 6b,c).

After transfection of the mutant library and selection for cells that had undergone successful recombination, each pool of mutant-expressing cells was exposed to varying concentrations of apoptolidin A or ammocidin A (Fig. 3a). Cells were cultured in galactose-containing media to enforce OXPHOS dependence and select against loss of function mutants. After two rounds of selection, resistant mutants were expanded, and the integrated variants were amplified by PCR and subjected to deep sequencing. Allelic enrichment was consistent across biological replicates and no enrichment was seen in cells passaged without inhibitors (Extended Data Fig. 6d). Comparison of the variant abundance in the plasmid library to the variant abundance in successfully integrated cells allows for inference of variant fitness in the absence of any treatment. Notably, ATP5B (β subunit) I393, L394, and E398, which all make close contacts with ammocidin in the atomic model of the inhibited ATP synthase, were enriched for wildtype alleles, suggesting that mutations at these positions result in a loss of fitness (Extended Data Fig. 6f).

Comparing variant abundance between the parental cell line and those treated with ammocidin A revealed two key mutational hot-spots: ATP5B-I390 and ATP5C-L77. Mutations at these positions conferred varying degrees of resistance to both apoptolidin and ammocidin (Fig. 3b, Extended Data Fig. 7), with positively charged residues exhibiting enrichment upon exposure to ammocidin A or apoptolidin A (Extended Data Fig. 6g,h). Substituting these mutations in the atomic model suggests that introduction of arginine at these positions likely alters binding site electrostatics and disrupts the hydrophobic interactions necessary for binding (Fig. 3d). Resistance was confirmed by expressing each mutation independently in 239LP cells (Extended Data Fig. 8a-c), as well as with CRISPR/Cas9 genome editing of the native alleles in K562 (Fig. 3c) and MV-4-11 leukemia cell lines (Fig. Extended Data Fig. 8k). Consistent with the results of the deep mutational scanning experiment, introduction of the ATP5B-I390R or ATP5C-L77R mutations conferred complete resistance to apoptolidin A and ammocidin A, while the ATP5B-I390Y variant raised the EC50s by 3 to 10-fold (Fig. 3c). None of the mutations tested notably affected sensitivity to oligomycin or puromycin, reflecting retention of a functional electron transport chain. These data establish that ATP synthase is the sole mechanistic target of apoptolidin family glycomacrolides; binding to the F_1_ subcomplex of ATP synthase is necessary for the selective cytotoxicity of the family.

**Fig. 3.**
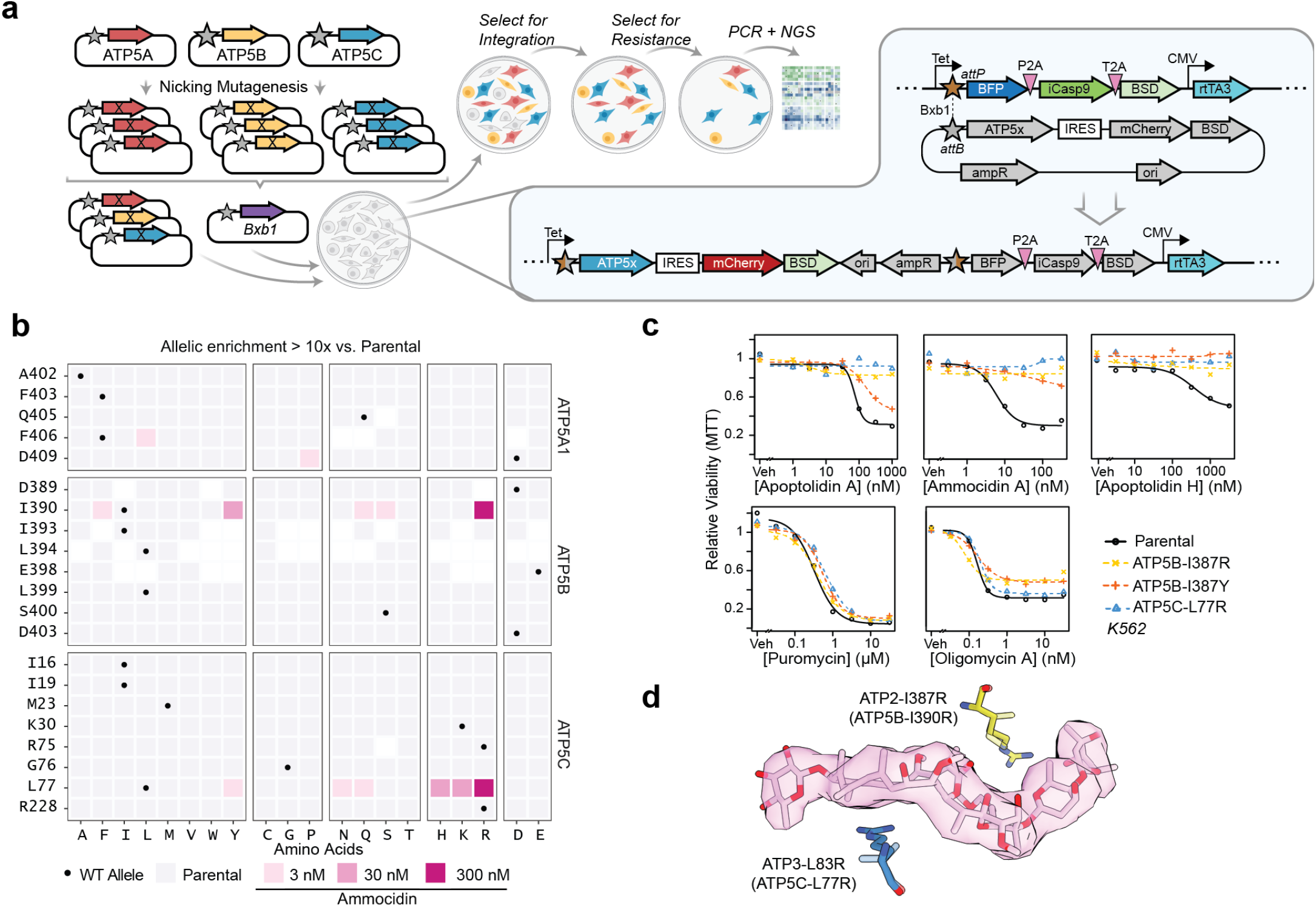
Deep mutational scanning and CRISPR/Cas9 directed editing of binding site residues reveals mutations which confer resistance to ammocidin and apoptolidin. **a,** Illustration of the Bxb1 landing pad system consisting of an *attP* containing landing pad integrated at a single site in the genome and an *attB* transfer plasmid containing the variant gene of interest (ATP5x), colored genes are expressed in the presence of doxycycline; **b,** Tile plot of allelic enrichment after selection, filled based on the highest dose at which >10-fold enrichment was observed. **c,** Comparison of drug sensitivity between K562 Parental and K562 CRISPR/Cas9 knock-in resistant mutants demonstrating cross resistance to glycomacrolides with retained sensitivity to oligomycin and puromycin; **d,** Modeling of L83R and I387R mutations in the ammocidin cryoEM model (~ human residues in parenthesis).

The binding modes of apoptolidin family macrolides revealed by cryoEM and deep mutational scanning studies also help contextualize the previous studies on the SAR of apoptolidin, which were notable in that no single modification was able to eliminate its activity^25,26^. This observation is consistent with its binding site, which is dominated by hydrophobic interactions each of which contributes a fractional amount to its nanomolar affinity. The structure also provides clues to the role of the disaccharide that is key for its activity: although loss of the disaccharide on apoptolidin H reduces its activity by >10 fold, it still retains sub micro-molar activity *ex vivo*. Importantly, this residual activity is eliminated by introduction of the apoptolidin resistance mutations ATP5B-I390R and ATP5C-L77R (Fig. 3c), suggestive of a shared binding mode of the macrolide core.

### Ammocidin A inhibits leukemia growth *in vivo*

Although apoptolidin and ammocidin are distinguished by their ability to induce apoptosis in transformed cells while sparing healthy cells, ATP synthase, the target of apoptolidin, is ubiquitously expressed and considered ‘essential’ in non-cancerous cells. Though the ATP synthase inhibitor oligomycin is widely used as a tool compound in cellular assays, its lipophilicity and narrow therapeutic index render it poorly suited for *in vivo* applications^47^. These studies on older ATP synthase inhibitors prompted us to question whether inhibition of ATP synthase by apoptolidin family glycomacrolides could be leveraged therapeutically and whether a therapeutic index between cancerous cells and healthy cells exists. Ammocidin A was selected as the lead compound for *in vivo* studies due to its serum stability, higher potency, and the known proclivity of apoptolidin A to isomerize *in vivo* to isoapoptolidin A via ester migration of C-20^48,49^. Preliminary pharmacokinetic studies confirmed that ammocidin injected intraperitoneally was bioavailable in blood (Extended Data Fig 9a) and dose escalation studies established ammocidin 0.1 mg/kg/day as the maximum tolerated dose, with minimal detectable toxicity over 14 days.

Based on the efficacy observed for ammocidin *in vitro*, we tested the compound as a monotherapy in an *in vivo* model. Antileukemic efficacy was evaluated in a systemic murine NSGS xenograft model^50^ using MV-4-11 human leukemia cells. Engrafted mice were treated with vehicle, 0.1 mg/kg or 0.03 mg/kg ammocidin A dosed daily, five days on, two days off, for two weeks (Fig. 4a). Weekly chimerism analyses were conducted, and the percentage of MV-4-11 cells were quantified via flow cytometric analysis of murine peripheral blood using anti-human CD45 (hCD45) and anti-human CD33 (hCD33) monoclonal antibodies. At day 28, vehicle mice became moribund, and all mice were sacrificed. Blood, bone marrow, and splenic tissues were harvested for chimerism analysis^50^. Treatment with ammocidin resulted in dramatically decreased bone marrow leukemia burden in a dose-dependent manner in mice treated with 0.1 mg/kg ammocidin (Fig. 4b). A trend towards leukemia suppression was noted in the spleen, however, this effect was not statistically significant. Leukemia burden was also assessed by immunohistochemistry for hCD45 in bone marrow, which similarly revealed dose-dependent leukemia suppression in ammocidin A treated mice (Fig. 4c). The remarkable degree of efficacy seen for ammocidin A monotherapy in leukemia is consistent with recent studies that suggest that leukemia cells, and in particular therapy resistant leukemia stem cells (LSCs), are exquisitely sensitive to OXPHOS inhibition^8,51–55^.

**Fig. 4.**
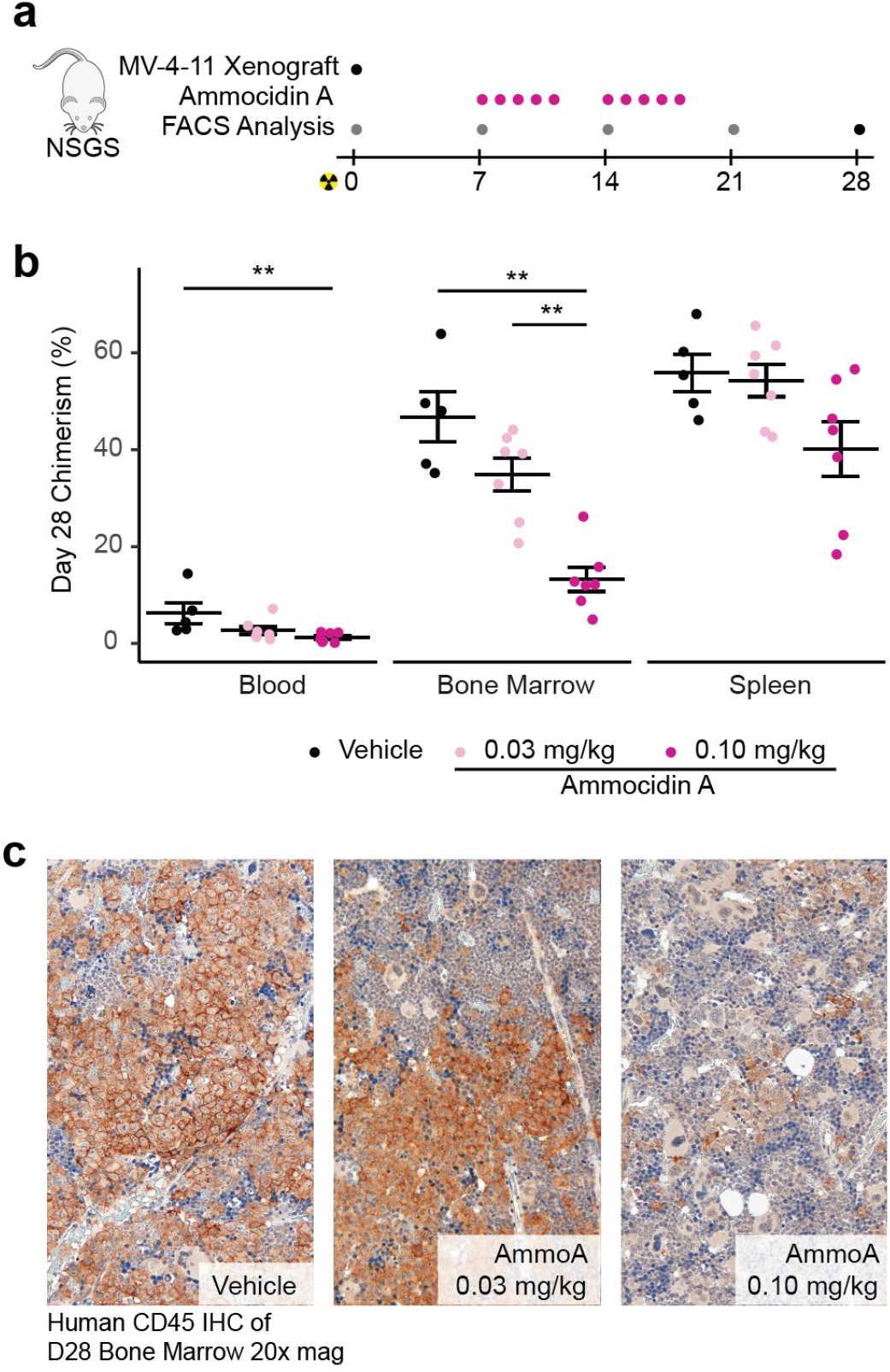
Ammocidin A inhibits leukemia growth in vivo. **a,** Experimental scheme for xenograft experiments—grey dots indicate assessment of human MV-4-11 chimerism in blood only, pink dots on days of ammocidin dosing, and black dot for terminal analysis on day 28 (e.g. assessment of chimerism in the blood, marrow, and spleen); **b,** Assessment of human MV-4-11 chimerism in the blood, bone marrow, and spleen at day 28; bars represent mean ± S.E.M for each group (n = 6 – 7 per group, ** = P < 0.01); **c,** Immunohistochemistry against human CD45 in day 28 marrow showing dose dependent decrease in leukemia burden.

## Discussion

Alterations in bioenergetics, including OXPHOS dependency, have long been recognized as a hallmark of cancer, however, efforts to target this vulnerability have been hampered by a dearth of metabolic inhibitors with sufficient selectivity and acceptable toxicity profiles^56^. This unmet need is especially acute as many of the cancers that appear to be most vulnerable to OXPHOS inhibition are those for which few therapeutic options exist, including acute myeloid leukemia and glioblastoma. Previous efforts to target mitochondrial bioenergetics in leukemia include inhibitors of mitochondrial complex I^51,57^, complex III^7^, and the mitochondrial translation machinery responsible for synthesizing the mitochondrially encoded components of the electron transport chain^58^. Outside of leukemia, therapeutic targeting of ATP synthase has previously been attempted in the context of autoimmune conditions with the OSCP binding benzodiazepine Bz-423^59^ which advanced to Phase II clinical trials, as well as the FDA approved anti-tuberculosis agent bedaquiline which targets the mycobacterial ATP synthase^60^. In addition to efforts to target the electron transport chain directly, recent studies have suggested that venetoclax, an inhibitor of BCL-2 recently approved for treating myeloid leukemias, targets LSCs by inhibiting OXPHOS^10,11,53^. There is also evidence that OXPHOS inhibitors can be combined synergistically to increase the efficacy of venetoclax therapy, suggesting a possible route for efficacious combination therapies with apoptolidin family glycomacrolides^58,61,62^. This work demonstrates that apoptolidin family glycomacrolides are unique among all OXPHOS inhibitors currently in clinical development or any previously described ATP synthase inhibitors.

Chemical probes, particularly compounds derived from natural products discovered in phenotypic screens^63^, have contributed much to our understanding about the functional dependencies of cancer cells. Though tool compounds may modulate pathways and phenotypes in desirable ways, to be considered a chemical probe a compound must act selectively through a defined mechanism of action^64^. In order for a compound to be considered a therapeutic candidate, it must be bioavailable, capable of modulating its target *in vivo,* and possess an acceptable therapeutic index^64^. Though ATP synthase was proposed as the target of apoptolidin based its cytotoxicity profile in the NCI-60 screen^18^, a lack of target validation precluded its use as a chemical probe and apoptolidin’s reduced stability precluded its development as a therapeutic agent. This work defines the mechanism of action of the entire class of apoptolidin glycomacrolides including apoptolidin A and ammocidin A, combining evidence from cellular target engagement, structural biology, and mutagenesis of the binding site. The *in vivo* studies using ammocidin A represent the first application of this compound class *in vivo* and demonstrate that ammocidin family compounds are bioavailable, well tolerated, and efficacious at suppressing leukemia growth in human xenografts. Combined with the structural insights and past work on the SAR of apoptolidin family compounds, this work establishes a path towards the therapeutic development of apoptolidin family glycomacrolides to address OXPHOS dependency in cancer and to study the reliance of cellular OXPHOS in human physiology and disease.

## Acknowledgements

We are grateful to Kenneth Matreyek, PhD of Case Western Reserve University for providing the Bxb1 landing pad cells and plasmids for the mutagenesis experiments as well as Ayesha Muhammad and Andrew Glazer, PhD for assistance applying the landing pad system. We are grateful to David Mueller, PhD of Rosalind Franklin University for providing the USY006 yeast strain and protocols for handling isolated mitochondria, as well as Pankaj Sharma, PhD for his assistance isolating F1 ATPase for enzymatic assays. We thank Kerry Brown, PhD and Hye-Young Kim, PhD for their helpful advice regarding synthesis and affinity enrichment. The ammocidin producing strain AJ9571 was provided by Ajinomoto Co., Inc., Tokyo, Japan under a material transfer agreement.

## Author contributions

B.J.R, G.A.S, L.P. and B.O.B. conceived of the project. B.J.R, H.G., H.E.R., M.T.W., B.I.R., and P.B.F designed the experiments. B.J.R. isolated compounds, sythesized and validated probes, performed affinity enrichments and gel based-profiling, and carried out mutational scanning and resistance experiements. H.G. prepared cryoEM grids, collected, and analyzed sturctural data. H.E.R performed xeograft experiments. M.T.W. carried out and analyzed proteomics experiments. B.I.R. generated stable cells lines and conducted toxicity studies. All authors contributed to the data analysis. B.J.R. and H.G. wrote the manuscript. P.B.F., W.K.R., L.P., M.R.S., J.L.R., and B.O.B. supervised the project.

## Data and materials availability

The atomic coordinates have been deposited in the Protein Data Bank (PDB) with the accession codes 7MD2 and 7MD3. The EM maps have been deposited in the Electron Microscopy Data Bank (EMDB) with the accession codes 23763, 23764 and 23765. The sequencing data and variant counts for the deep mutational scanning experiments have been depositing in the Gene Expression Omnibus (GEO) database under accession code GSE171362. Materials and compounds are available from the corresponding author on request.

## Materials and Methods

### Cell Culture

MV-4-11, HEK-293T, and H-292 cells were purchased from the American Type Culture Collection (ATCC, Manassas, VA). K562 cells were kindly provided by Dr. Gregor Neuert respectively and were validated by STR profiling at the University of Arizona genetics core. BaF3 cells were a kind gift of Dr. Christine Lovly, Vanderbilt University Medical Center. HEK-293-LP cells were a kind gift of Kenneth Matreyek, Case Western Reserve University. Cell lines were tested for mycoplasma contamination using the ATCC Universal Mycoplasma detection kit before use. MV-4-11 and K562 cells were maintained with Gibco IMDM Glutamax media supplemented with 10% FBS and 1% Pen/Strep. H292 cells were maintained with RPMI 1640 supplemented with 10% FBS and 1% Pen/Strep. HEK-293 cells were cultured in DMEM media, 10% FBS, 1% Pen/Strep. Cells grown with galactose media were grown in glucose free DMEM media supplemented with 10% FBS, 1% Pen/Strep, and 10 mM galactose. Cells were maintained at a density of 5 ×10^4^− 1 × 10^6^, except where single cells were isolated to generate clonal populations.

### Molecular Biology

All plasmids were prepared using commercial MiniPrep kits (Qiagen, Hilden, Germany). All enzymes were obtained from New England Biolabs and used per the manufacturer’s recommendations. PCR reactions were carried out using Q5 High-Fidelity Polymerase (New England Biolabs, Ipswich, MA) per the manufacturer’s protocols. Primers were ordered from Genosys (Sigma-Aldirch, Burlington, MA) or Integrated DNA Technologies (Coralville, IA). Except where noted, genomic DNA was extracted using QuikExtract DNA reagent (Lucigen, Middleton, WI). Except where noted, non-viral plasmids were maintained in competent DH5-alpha *E. coli*, prepared by the Vanderbilt Molecular Cell Biology Resource (MCBR), lentiviral and retroviral plasmids were maintained in Stbl2 *E. coli* (ThermoFisher, Waltham, MA), and cloned plasmids were initially transformed into competent cells included with kits. Gibson assemblies were carried out using the NEB Gibson Assembly Cloning kit. Site Directed Mutagenesis was carried out using the QuikChange Lightning II cloning kit (Agilent, Santa Clara, CA). PCR cloning was carried out using the NEB PCR cloning kit. All Sanger sequencing was completed by Genewiz, Inc (South Plainfield, NJ).

### Immunoblotting

Cell lysates were prepared using RIPA buffer (see lysis and click chemistry methods below) and run on SDS Tris-Glycine gels (Precast 4 - 20% Bio-Rad, or hand-casted 20%). Gels were transferred to low-background PVDF membranes using the BioRad TransBlot Turbo system and blocked with 5% milk powder in TBS + 0.1% Tween 20 (TBST) for 1 hr at room temperature. Primary antibodies were diluted at 1:1000 unless otherwise specified in 5% BSA in TBST + 0.1% sodium azide and incubated overnight at 4 °C. Membranes were washed for 5 min × 5 with TBST, and appropriate Starbright Blue 700 secondary antibody (Bio-Rad, Hercules, CA) for 1 hr at room temperature, followed by 5 × 5 min washes with TBST. Membranes were imaged using the ChemiDoc MP imaging system. Antibodies used for immunoblotting were: ATP5B (Proteintech, 17247–1AP), VCAC1 (CST #4661), TIMM17B (Proteintech 11062–1-AP), FLAG^®^ (Sigma Aldrich, M2), ATP5G1 (Abcam, EPR13908), Actin (BioRad, 12004163).

### Photoaffinity Labeling

Photoaffinity labeling was carried out essentially as described in Thomas JR, 2017^65^. Briefly, ~5 - 30M cells per condition were washed twice with OptiMEM media and resuspended in 1 mL of OptiMEM media in a 12 well plate. Each well was treated with vehicle (DMSO) or competitor as indicated for 1 hr at 37 °C followed by the addition of vehicle or photoaffinity probe for 1 hr at 37 °C. The samples there then irradiated with 365 nm light for 15 min on ice using a Stratalinker 2400 (Agilent). The samples were then washed twice with 1 mL of 1x PBS and then lysed as described below or flash frozen at −80 °C for later processing.

### Lysis and Click Reaction

Samples were processed essential as described in Paxman et al, 2018 ^66^ with minor modifications. Cell pellets were lysed in 100 μL of radioimmunoprecipitation assay buffer (RIPA: 150 mM NaCl, 50 mM Tris pH 7.5, 1 % Triton X-100, 0.5% sodium deoxycholate, 0.1% SDS). After lysis, the samples were centrifuged for 1 min at 28,000 g and the lysates transferred to a fresh tube. Protein concentration was determined using the BCA assay (Pierce) and each sample was adjusted to a concentration of 1 mg/mL. Each 100 - 1,000 μg click reaction was carried out at the following final concentrations: 100 μM azide (as indicated: Azide Fluor 545 – Sigma Aldrich OR TAMRA-Azide-Desthiobiotin – Broadpharm), 800 μM CuSO_4_, 1.6 mM BTTAA ligand (Click Chemistry Tools), and 5 mM sodium ascorbate. The azide was added directly to the cell lysate while the CuSO_4_, BTTA, and sodium ascorbate (fresh) were prepared as a master mix and added to each tube. The samples were vortexed briefly and incubated at 37 °C for 1 hr. The proteins were precipitated using a methanol-chloroform precipitation using a 3:1:4 ratio of methanol, chloroform, and water. The pellet was washed once with 3:1 MeOH/CHCl_3_, and once with methanol and dried briefly at room temp, taking care to avoid over drying the pellet. Samples used for in-gel analysis were resuspended in 1 × loading buffer + BME (Bio-rad), heated for 15 min at 37 °C followed by sonication for 15 min to aid solubilization. The samples were boiled for 5 min at 95 °C, centrifuged at 20,800 g × 1 min, and 10 μg of protein was loaded into each lane. Samples used for affinity purification were processed as described below.

### Affinity Enrichment

Protein pellets functionalized with the TAMRA-Azide-Desthiobiotin (Click Chemistry Tools, Scottsdale, AZ) were precipitated by methanol/chloroform as described above and resuspended in 500 μL of 6M Urea + 25 mM ammonium bicarbonate, with 140 μL of 10% SDS was added to aid resolubilization. 6 mL of 1 x PBS was added to each sample to lower the urea concentration to < 0.5 M. 50 μL of high-capacity streptavidin beads (Pierce, Waltham, MA) were washed three times with PBS, added to each sample, and incubated for 2 hrs at room temperature on a rotator. Each sample was then loaded onto a 0.8 mL spin column (Pierce) in 500 μL portions using a vacuum manifold. The beads were then washed with 4 × 500 μL volumes of 1% SDS in PBS, 4 volumes of 4M Urea in PBS, 4 volumes of 1M sodium chloride, and 1 volume of 1% SDS in PBS. The bead slurry was then resuspended in 100 μL of 50 mM biotin, 1% SDS, in PBS (pH 7.2), incubated at 37 °C for 10 minutes, agitated for 10 minutes, and then collected into a fresh 1.5 mL Eppendorf tube via centrifugation at 5200 g x 3 min. The biotin elution was repeated once with a second 100 μL volume of 50 mM biotin, 1% SDS in PBS. The two biotin elution fractions were combined and precipitated with methanol/chloroform as described above and resuspended in either 1 x loading buffer (if running a gel and silver staining) or in 0.1% RapiGest SF (Waters Corp, Milford, MA) in 0.1 M HEPES pH 8.0 if proceeding to MudPIT proteomics. Silver staining was carried out using the Pierce Silver Stain kit, per the manufacturers protocols.

### TMT-Multiplexed Proteomics Sample Preparation

TMT samples were prepared as follows: RapiGest resuspended samples (in 50 μL) were reduced by the addition of 0.5 μL 0.5 M TCEP (final concentration, 5 mM) for 30 min at room temperature and cysteines alkylated using 1 μL 0.5 M iodoacetamide (final concentration, 10 mM) for 30 min at room temperature protected from light. Samples were digested by addition of 0.25 μg of sequencing grade trypsin dissolved in 50 mM acetic acid at 0.5 μg/mL (Promega) overnight at 37 °C with vigorous shaking. Sample volume was adjusted to 60 μL with H_2_O and TMT labeling was carried out using the TMT sixplex isobaric label (Thermo Scientific) using 100 μg of reagent dissolved in 100% acetonitrile per sample and incubated at room temperature for 1 hr. The labeling reaction was quenched with 4 μL of 10% w/v of ammonium bicarbonate (final concentration of 0.4% w/v) at room temperature for 1 hr. Samples were then acidified with formic acid (final conc 5% v/v) and concentrated *in vacuo* to ~1/6^th^ of its original volume to remove the acetonitrile and sample volume was adjusted to 600 μL with 0.1% formic acid. Samples were then heated at 42 °C for 30 min to precipitate the RapiGest surfactant and insoluble components were removed by centrifugation at 28,000 g x 30 min. Digested, TMT-labeled samples were stored at −80 °C until analysis.

### In-gel Digest Proteomics Sample Preparation

Gel bands were excised and washed three times with 25 mM ammonium bicarbonate in 1:1 acetonitrile/water, and then dried using a vacuum centrifuge. Proteins were reduced with the addition of 10 mM DTT in 25 mM ammonium bicarbonate in 10% acetonitrile (v/v), and heating at 56 °C for 1 hr. Next, proteins were alkylated with 55 mM iodoacetamide at room temperature for 45 minutes. Gel pieces were then rinsed with 25 mM ammonium bicarbonate is aqueous solution, followed by 25 mM ammonium bicarbonate in 1:1 acetonitrile/water. Rinses were repeated and gels pieces were dried using a vacuum centrifuge. Enough trypsin (12.5 ng/μL in 25 mM ammonium bicarbonate in aqueous solution) was added to cover the gel pieces and samples were incubated on ice for 30 min. Excess trypsin solution was removed and 25 mM ammonium bicarbonate in aqueous solution was added. Samples were then incubated at 37 °C for 8 hrs. After digestion, three volumes of water were added to the gel samples and they were vortexed for 10 min, sonicated for 5 min, and the resulting supernatant was collected. Gel pieces then had 45% water/50% acetonitrile/5% formic acid (v/v/v) added and were vortexed for 10 min, sonicated for 5 min, and the resulting supernatant was collected. The extraction with 45% water/50% acetonitrile/5% formic acid (v/v/v) was repeated and supernatant was again collected. Samples were concentrated using a vacuum centrifuge and analyzed.

### Liquid Chromatography – Tandem Mass Spectrometry

MudPIT microcolumns were prepared and peptide samples were directly loaded onto the columns using a high-pressure chamber. Samples were then desalted for 30 min with buffer A (95% water, 4.9% acetonitrile, 0.1% formic acid v/v/v). LC-MS/MS analysis was performed using a Q-Exactive HF (Thermo Fisher) mass spectrometer equipped with an Ultimate-3000 RSLCnano system (Thermo Fisher). MudPIT experiments were performed with 10 μL sequential injections of 0, 10, 30, 60, and 100% buffer C (500mM ammonium acetate in buffer A), followed by a final injection of 90% buffer C with 10% buffer B (99.9% acetonitrile, 0.1% formic acid v/v) and each step followed by a 130 minute gradient from 5% to 80% B with a flow rate of 300 nL/min on a 20 cm fused silica microcapillary column (ID 100 μm) ending with a laser-pulled tip filled with Aqua C18, 3 μm, 125 Å resin (Phenomenex). Electrospray ionization (ESI) was performed directly from the analytical column by applying a voltage of 2.0 V with an inlet capillary temperature of 275 °C. Data-dependent acquisition of mass spectra was carried out by performing a full scan from 300–1800 m/z with a resolution of 60,000. The top 15 peaks for each full scan were fragmented by HCD using normalized collision energy of 38, 0.7 m/z isolation window, 120 ms maximum injection time, at a resolution of 15,000 scanned from 100 to 1800 m/z and dynamic exclusion set to 60 s. Peptide identification and TMT-based protein quantification was carried out using Proteome Discoverer 2.3. MS/MS spectra were extracted from Thermo Xcalibur .raw file format and searched using SEQUEST against a Uniprot human proteome database (released 03/2014 and containing 20337 entries). The database was curated to remove redundant protein and splice-isoforms and supplemented with common biological MS contaminants. Searches were carried out using a decoy database of reversed peptide sequences and the following parameters: 10 ppm peptide precursor tolerance, 0.02 Da fragment mass tolerance, minimum peptide length of 6 amino acids, trypsin cleavage with a maximum of two missed cleavages, dynamic methionine modification of 15.9949 Da (oxidation), static cysteine modification of 57.0215 Da (carbamidomethylation), and static N-terminal and lysine modifications of 229.1629 Da (TMT sixplex).

### Statistical Analysis of TMT Proteomics

DataProtein matches and TMT quantification were processed using a custom R script using the DEP pipeline ^67^. Briefly, raw TMT quantifications were normalized using the vsn package ^68^. Differential enrichment was determined using limma ^69^. Enriched proteins were selected based on an adjusted p-value of 0.05 and log2 fold change of 2 (see Data S2). Plots were generated using ggplot2.

### MTT Viability Assay

The MTT assay protocol was adapted from Deguire *et al.* ^*32*^. Briefly, a 100 μL of suspension cells at 100,000 per mL were added to wells of a microtiter plate precoated with 0.5 μL of test compound at 200x in DMSO and incubated for 48 - 72 hrs (as noted). MTT reagent was dissolved in fresh media at 1 mg/mL and 100 μL was added to each well to achieve a final concentration of 0.5 mg/mL and incubated for 2 hrs at 37 °C. Cells were centrifuged at 800 g x 5 min and decanted. MTT crystals were redissolved in 100 μL of DMSO, allowed to incubate for 5 min at RT, and read at 560 nM using a SpectraMax plus 384 plate reader (Molecular Devices, San Jose, CA). Absorbance values were normalized by background subtraction (wells without cells) and normalized such that vehicle treated cells had a viability of 1.0. IC_50_ curves were fit using a custom R script using the DRC package ^70^ with a four-parameter log-logistic function.

### Sulforhodamine B (SRB) Proliferation Assay

The SRB assay protocol was adapted from ^71^ to assess cytotoxicity in adherent cells (H292 and HEK-293). Cells were treated as above in the MTT assay in TC treated 96 well plates with 100 μL of media. After 48 – 72 hrs, cells were fixed by the gentle addition of 25 μL of cold 50% (w/v) trichloroacetic acid (TCA) and incubated at 4 °C for 1 h. Plates were then washed 4x by submersion in a basin of tap water and allowed to dry overnight. Cells were stained by the addition of 50 μL of 0.04% (w/v) SRB dissolved in 1% (v/v) acetic acid to each well and allowed to incubate at room temperature for 1 hr. The SRB solution was decanted and plates were washed with 4x 100 μL volumes of 1% (v/v) acetic acid and allowed to dry. The SRB reagent was redissolved by the addition of 10 mM Tris base (pH 10.5), allowed to incubate for 10 min, and absorbance was measured at 510 nM using a SpectraMax plus 384 plate reader (Molecular Devices, San Jose, CA). Absorbance values were normalized by background subtraction (wells without cells) and normalized such that vehicle treated cells had a viability of 1.0. IC_50_ curves were fit using a custom R script using the DRC package ^70^ with a four-parameter log-logistic function.

### Confocal Microscopy

Coverslips, (22 mm diameter, 1.5H thickness) (Thorlabs) were washed with 1M HCl for 1 hrs at 55 °C, washed with DI water and 70% ethanol, and coated with poly-D-lysine (100 μg/mL in sterile water), and allowed dry. Adherent cells were plated on top of coverslips and grown till confluent. For imaging of apoptolidin A PA localization, cells were treated with 200 μM apoptolidin A PA for 1 h and photocrosslinked for 15 min as described under ‘photoaffinity labeling’. Mitotracker Deep Red FM (Invitrogen) was added 30 min prior to the end of the experiment at a final concentration of 100 nM. Cells were washed 2x with PBS (containing Ca^++^ and Mg^++^) and fixed with 1.6% PFA for 15 min. Apoptolidin A PA treated cells were permeabilized with ice cold methanol for 20 min at −20 °C, while cells intended for immunofluorescence were permeabilized with 1% Triton X100 for 15 min. Cells were washed with 1% BSA in PBS twice, while cells intended for antibody staining were then blocked with 5% BSA in PBS. Azide Fluor 545 labeling was carried out as described under ‘lysis and click reaction’ with the azide concentration reduced to 100 μM, and samples were washed with 1% BSA in PBS after 1 hr at 37 °C. Samples for immunofluorescence were stained for overnight with primary antibodies, washed 4x with 1% BSA in PBS, followed by 1 hr staining at room temperature with secondary antibodies (goat anti-mouse alexa 488 antibody, Invitrogen A-11008), followed by 4x washes with 1% BSA in PBS. Samples were stained briefly with DAPI (1 μg/mL) mounted on slides using ProLong Glass mountant (Invitrogen), allowed to set for 72 hrs and imaged using a Zeiss 880 Airyscan confocal microscope.

### Yeast growth and ATP synthase purification

ATP synthase was purified from yeast strain USY006 ^72^ bearing 6×His tags at the N-termini of the β subunits as described previously but with the following modifications^73^. Yeast was cultured in an 11 L fermenter (New Brunswick Scientific) and the mitochondria were prepared by breaking yeast cell wall by bead beating. All subsequent purification steps were performed at 4 °C. Mitochondria were washed with phosphate buffer (50 mM sodium phosphate, pH 9.0, 5 mM 6-aminocaproic acid, 5 mM benzamidine, 1 mM PMSF) for 30 min before being collected by centrifugation at 184,000 g for 30 min. Membranes were solubilized by resuspending in buffer (50 mM Tris-HCl, pH 7.4, 10% [v/v] glycerol, 1% [w/w] dodecyl-β-D-maltoside [DDM, Anatrace], 5 mM 6-aminocaproic acid, 5 mM benzamidine, 1 mM PMSF) and mixed for 1 hour. Insoluble material was removed by centrifugation at 184,000 g for 30 min and the supernatant containing solubilized protein was collected. Imidazole was added to 40 mM and NaCl to 300 mM and the sample was loaded onto a HisTrap HP 5 mL column (MilliporeSigma, Burlington, MA) equilibrated with wash buffer (50 mM Tris-HCl, pH 7.4, 10% [v/v] glycerol, 0.05% [w/w], 40 mM imidazole, 300 mM NaCl, 5 mM 6-aminocaproic acid, 5 mM benzamidine, 1 mM PMSF). The column was washed with 5 column volumes of wash buffer and ATP synthase was eluted with the wash buffer containing 300 mM imidazole before being loaded to a Superose 6 increase column (MilliporeSigma) equilibrated with buffer (20 mM Tris-HCl, pH 7.4, 10% [v/v] glycerol, 0.05% [w/w], 100 mM NaCl, 5mM MgCl_2_). Fractions containing ATP synthase were pooled and protein was concentrated to ~15 mg/ml prior to storage at −80 °C.

### Yeast F_1_-ATPase Preparation

Preparation of F1-ATPase followed the same protocol as intact ATP synthase preparation until the mitochondrial were pelleted after bead beating. Protocols were derived from Mueller, 2004 ^74^. Unless otherwise noted, all steps were done at 4 °C. Submitochondrial particles (SMPs) were prepared by resuspending mitochondria at 10 mg/mL in sonication buffer (SB - 0.25 M sucrose, 50 mM phosphate buffer, 5 mM 6-aminocaproic acid, 5 mM benzamidine, 1 mM PMSF, pH 7.5) with 1 mM EDTA and sonicated at 150W × 1 min in 5 s on 10 s off cycles, and centrifuged at 5000 g for 10 min. The supernatant containing SMPs was centrifuged at 100,000 g for 60 min at 4 °C. The pellet was washed twice with SB without EDTA, resuspended at 20 mg/mL, and warmed to room temperature. F1 ATPase was extracted from SMPs by the addition of 0.5 volumes of PBS saturated chloroform and vortexed for 30s. ADP was added at a final concentration of 2 mM and the phases were separated by centrifugation at 3000 g for 10 min at 15 °C. The aqueous layer was collected, methanol was added to a final concentration of 10% v/v, followed by the addition of Buffer D (10% MeOH, 1.2 M NaCl, 40% [w/v] glycerol, 0.25 M sucrose, 50 mM phosphate, 5 mM 6-aminocaproic acid, 5 mM benzamidine, pH 7.5). The crude F1 preparation was loaded onto a 5mL HisTrap HP 5 mL column at 4 °C and washed with 300 mL of 97 % wash buffer (10% MeOH, 10% glycerol, 0.3M NaCl, 0.25 M sucrose, 50 mM phosphate, 5 mM 6-aminocaproic acid, 5 mM benzamidine, 1 mM PMSF, pH 7.5) + 3% elution buffer (wash buffer + 0.4 M imidazole). F1 ATPase was eluted with 100% wash buffer and the F1-containing fractions were pooled and concentrated to < 1 mL. F1 ATPase was further purified by means of gel-filtration at room temperature with a Superdex 200 increase 10/300 GL column with SDX buffer (0.25 M sucrose, 0.2 M NaCl, 50 mM Tris, 1 mM EDTA, 1 mM ATP, 0.5 mM PMSF, pH 8.0) at 0.5 mL/min, the F1 containing fractions were pooled, and precipitated by the addition of saturated ammonium sulfate (70%), and stored at 4 °C.

### CryoEM Grid Preparation

CryoEM grids of drug-free ATP synthase were prepared by removing glycerol from the sample with Zeba Spin Desalting Column (Thermo Fisher Scientific) immediately before freezing. Glycerol-free ATP synthase was then applied to home-made ^75^ holey-gold grids ^76,77^ that had been glow discharged in air for 2 min. Grids were blotted with modified Vitrobot Mark III (Thermo Fisher Scientific) for 26 s with ~100% humidity at 4 °C before being plunge frozen in a liquid ethane/propane mixture ^78^. Grids of ATP synthase with ammocidin and apoptolidin were prepared the same way except that ATP synthase was incubated with either 60 μM ammocidin or 20 μM apoptolidin for 1 hour and the glycerol removal step was performed with buffers containing the same concentrations of inhibitors.

### CryoEM Data collection

CryoEM movies of yeast ATP synthase bound to ammocidin and apoptolidin were collected with a Titan Krios G3 electron microscope operated at 300 kV and equipped with a prototype Falcon 4 camera (Thermo Fisher Scientific). Automatic data collection was performed with *EPU* (Thermo Fisher Scientific). For the ammocidin dataset, 4345 movies each containing 30 fractions were collected at a nominal magnification of 75000×, corresponding to a calibrated pixel size of 1.03 Å. The total exposure and the camera exposure rate were ~45 e^−^/Å^2^ and 5.0 e^−^/pix/s, respectively. The apoptolidin dataset, consisting of 4019 movies with 29 fractions each were collected at same magnification as the ammocidin dataset. The total exposure and the camera exposure rate for this dataset were ~43 e^−^/Å^2^ and 4.8 e^−^/pix/s, respectively. The drug-free ATP synthase dataset was collected on a Tecnai F20 electron microscope (Thermo Fisher Scientific) operated at 200 kV and equipped with a K2 Summit camera (Gatan). A dataset consisting 190 movies each containing 30 fractions was manually collected at a nominal magnification of 25000×, corresponding to a calibrated pixel size of 1.45 Å. The total exposure and the camera exposure rate were ~36 e^−^/Å^2^ and 5.0 e^−^/pix/s, respectively.

### Image analysis

Image analysis was performed with *cryoSPARC v2* ^79^ except where noted. Movie frames were aligned with *MotionCor2* ^*80*^ using a 7×7 grid and contrast transfer function (CTF) parameter were estimated in patches. Two templates displaying a sideview and an oblique view of ATP synthase for particle picking were generated with 2D classification of manually selected particles with a box size of 320×320. Template picking generated 654,097 particle images for the ammocidin dataset, 1,189,085 particle images for the apoptolidin dataset and 38,514 particle images for the drug-free dataset. After cleaning by 2D classification, 329,297, 621,136, and 34,035 particle images remained for the corresponding datasets. Ab initio 3D classification and heterogeneous refinement identified three classes for each dataset corresponding to the three main rotational states of ATP synthase. For each dataset, the maps of the three states were aligned based on the F1 region and a masked local non-uniform refinement ^81^ of the region was performed. For the high-resolution ammocidin and apoptolidin datasets, individual particle defocus parameters were estimated, and a second masked refinement was performed with updated CTF parameters. Image parameters were then converted to *Relion 3.0* ^*82*^ format with the *pyem* package ^83^ and individual particle motion was re-estimated with Bayesian Polishing ^84^. For the apoptolidin dataset, particle images were downscaled to a pixel size of 1.2875 Å with *Relion 3.0*. The resulting particles were imported back to *cryoSPARC v2* and another round of masked non-uniform refinement of the F1 region was performed. For the ammocidin and apoptolidin datasets, individual particle defocus was re-estimated, followed by a final round of masked refinement. The final maps of the ammocidin, apoptolidin, and drug-free dataset contain 289,501, 477,847 and 34,035 particle images and reached resolutions of 3.1 Å, 3.3 Å and 4.2 Å, respectively. The ammocidin and apoptolidin maps were locally sharpened with a *COSMIC2* ^*85*^ implementation of *DeepEMhancer* ^*86*^.

### Atomic model building

To model the F_1_ region of the ammocidin-bound structure, individual subunits of the F_1_ region of a yeast ATP synthase crystal structure (2XOK) was fit rigidly into the ammocidin map with *UCSF Chimera* ^*87*^. The model was then manually adjusted in *Coot* ^*88*^ and *ISOLDE* ^*89*^ before being refined in *Phenix* ^*90*^. The model of the apoptolidin bound structure was built in a similar manner, with the ammocidin model as the starting structure. To demonstrate the conformational change induced upon inhibitor binding, individual subunits of the same crystal structure (2XOK) were fit into the drug-free map as rigid bodies to generate Figure 2D and Movie 1. Figures and supplementary movie were generated with *UCSF Chimera* and *USCF ChimeraX* ^91^.

### Isolation of Mouse Liver Mitochondria

Mouse liver mitochondria were isolated essentially as described by Frezza et al.^92^ All steps were performed on ice. Freshly harvested mouse livers with the gallbladder removed were washed four times in mitochondrial isolation buffer (IB = 0.2 M sucrose, 10 mM Tris-MOPS, 1 mM EGTA/Tris, pH 7.4), weighed, and cut into small pieces with scissors. Liver pieces were resuspended in 2 volumes of IB, transferred to a 45 mL Potter-Elvehjem homogenizer, and homogenized with four strokes of the pestle. The homogenate was diluted with an additional 3 volumes of IB, transferred to a 50 mL conical vial, and centrifuged at 600g × 10 min at 4 °C to pellet nuclei. The supernatant was transferred to a 50 mL Nalgene Oak Ridge Centrifuge Tube and centrifuged at 7000g × 10min at 4 °C in a Sorvall RC 2-B centrifuge equipped with an SA-600 rotor to pellet mitochondria. The mitochondrial pellet was resuspended in 5 volumes of IB and centrifuged at 7000g x 10 min at 4 °C a second time to remove microsomes. The mitochondrial pellet was resuspended at 20 mg/mL as determined using a BCA assay (Pierce).

### Enzyme Coupled ATP Synthesis Assay

The ATP synthesis assay was adapted from assays described by Gouspillou ^93^ and Cross ^94^. The respiration buffer (RB) consisted of 240 mM mannitol, 100 mM KCl, 1mM EGTA, 20 mM MgCl_2_, 10 mM KH_2_PO_4_, and 0.1% (w/v) fatty acid free BSA (pH 7.2). 20 mM Succinate, pH 7.2 with KOH (S3674) was used as a substrate in all experiments. The assay readout consisted of 5 mM glucose, 2.5 U/ml hexokinase (Sigma H4502), 2.5 U/ml glucose-6-phosphate dehydrogenase (Sigma G8529), and 1.6 mM NADP^+^(Sigma N8035). 20 μM P^1^,P^5^-di(adenosine-5) pentaphosphate (AP5A) was used to inhibit adenylate kinase which could otherwise convert 2 ADP → AMP + ATP. The assay reagents were prepared at 2 × in RB and 100 μL was dispensed into each well of UV transparent 96 well plate. 50 μL of 4x succinate was then added, followed by 25 μL of 8x mitochondria. The plate was incubated at 25 °C for 5 min to allow the mitochondria to energize. The reaction was initiated by the addition of 25 μL of 8x ADP using a multichannel pipette to yield a total volume in the well of 200 μL. Absorbance was monitored at 340 nM using a SpectraMax Plus 384 plate reader (Molecular Devices) set at 25 °C.

### Enzyme Coupled ATPase Assay for F_1_ ATPase

The ATPase assay was adapted from Cross ^94^. The assay buffer consisted of 60 mM Tris-Acetate pH 7.8; 1 mM MgCl_2_, 2.5 mM phosphoenolpyruvate; 1 mM KCN. The working solution was prepared at 2 × for a final concertation of 0.4 mM NADH; 3 U/mL of pyruvate kinase (PK) + 4.5 U/mL Lactate Dehydrogenase (LDH) [Sigma P0294]; ±10 μM carbonylcyanide p-trifluoromethoxyphenylhydrazone (FCCP). Unless otherwise specified, ATP was added as a 4x solution at 200 μM. Absorbance was monitored at 340 nm using a SpectraMax Plus 384 plate reader (Molecular Devices) set at 25 °C. 50 μL of enzyme was added to 100 μL of working solution and monitored for 2 min, 50 μL of ATP was then added for a final volume of 200 μL and absorbance was monitored over 10 minutes.

### Enzyme Coupled ATPase Assay for ATP synthase

To measure the ATP hydrolysis activity of ATP synthase, enzyme-coupled ATPase activity assays ^95^ were performed in 96-well plates with 160 μL buffer (50 mM Tris-HCl pH 7.4, 150 mM NaCl, 10% [v/v] glycerol, 5 mM MgCl_2_, 0.2 mM NADH, 2 mM ATP, 1 mM phosphoenol pyruvate, 3.2 units pyruvate kinase, 8 units lactate dehydrogenase, 0.05% [w/v] DDM, DMSO [v/v] 2%, 2 nM ATP synthase). NADH concentration was monitored at 24 °C **with a Synergy Neo2 Multi-Mode Assay Microplate Reader** (BioTek) measuring absorbance at 340 nm. Inhibition with ammocidin and apoptolidin was determined by adding different concentrations of the inhibitors to assay mixture. Assays were performed in triplicates with two independently purified batches of yeast ATP synthase.

### Preparation of Lentivirus

H293T were maintained in high glucose DMEM supplemented with 10% FBS, without antibiotics at approximately 50 - 60% confluence on the day of transfection. The transfection mixture consistent of 1 μg lentiviral plasmid, 750 ng psPAX2 (Addgene #12260), and 250 ng of pMD2.G (Addgene #12259) using FuGENE 6 per manufacturers protocols. The cells were incubated with plasmid for 16 hrs at which point the media was replaced with DMEM supplement with 10% FBS and 1% Pen/Strep. Viral supernatant was harvested at 24 hrs and 48 hrs and sterile filtered using 45 μM syringe filtered. Viral media was used immediately or frozen at −80 °C for later use.

### Lentiviral Transduction

MV-4-11 were seeded into 6 well plates in 2 mL of media. 500 μL of viral supernatant was added to each well, along with 500 μL of 36 μg/mL DEAE-Dextran in DMEM + 10% FBS + 1% Pen/Strep, for a final DEAE-Dextran concentration of 6 μg/mL. Cells were allowed to incubate with virus for 48 hrs, at which point antibiotics were applied for selection or cells were single cell sorted to generate clones.

### Retroviral Production

Platinum-E (Plat-E) cells selected for 2 days in DMEM + 10% FBS + 10 μg/mL blasticidin + 1 μg/mL puromycin in 100 mm tissue culture treated dishes. Selection media was replaced with DMEM + 10% FBS without antibiotics overnight. A transfection mix consisting of 500 μL OPTI-MEM + 30 μL FuGENE 6 was allowed to incubate for 5 min at room temperature. 10 μg of plasmid (pBABE-puro, Addgene #1764 or pBabe-puro Ras V12, Addgene #1768) was added to the FuGENE + OPTI-MEM, and allowed to incubate for 10 min. The media was removed and replaced with fresh DMEM + 10% FBS + 1% PenStrep. Viral supernatant was harvested after 48 hrs, sterile filtered, and stored at −80 °C.

### Retroviral Transduction

Retroviral transduction in BaF3s (gift of Dr. Christine Lovly) were conducted essentially as described in Gallant, 2015 ^96^. Cells were seeded at 50,000 /mL per well of a 6 well plate in RPMI 1640 + 10% FBS + 1% PenStrep, supplemented with recombinant murine IL-3 (Gibco, PMC0035) at 1 μg/mL (BaF3). 1 mL of viral supernatant was added to each well along with polybrene in DMEM for a final polybrene concentration of 10 μg/mL. The cells were allowed to incubate with virus for 48 hrs, at which point the viral media was removed and replaced with RPMI 1640 + 10% FBS + 1% PenStrep + IL-3 + 2 μg/mL Puromycin.

### ATP Synthase Mutant Library Preparation

ATP synthase ORF clones were purchased from GenScript USA Inc (Piscataway, NJ, USA) [ATP5A1, NM001001937.1; ATP5B NM_001686.4; ATP5C NM_001001973.3], PCR amplified, and cloned into the NotI digested pAG490 ^97^ using Gibson assembly and were confirmed by Sanger sequencing using ag122 and ag123 primers (See Data S2). Mutagenic primers were designed in three pools, one for each ORF, using the published primer design script, specifying a single codon for each of 20 amino acids without stop codons at a total of 21 positions (420 total variants), with 30 nt homology arms on both sides of the mutated codon (see Data S2 for primer sequences) ^98^. Primers were obtained as three separate 50 pm oPools from Integrated DNA technologies (Coralville, IA, USA). Nicking mutagenesis was carried out exactly as described in Wrenbeck EE, 2016 ^46^, Supplementary Protocol 1, except for reversing the use of the Nt.BbvCI and Nb.BbvCI enzymes to degrade the proper strands. Primer ag123 was used to synthesize the complementary strand. The product of each mutagenesis reaction was transformed into XL1-Blue Electroporation-Competent Cells (Agilent Technologies, Santa Clara, CA, USA) and serial dilutions were plated to determine the total number of colonies. All colonies from the 25 cm x 25 cm LB plate were collected, and plasmid was purified by MiniPrep. The three pools were combined in a 5:8:8 ATP5A1:ATP5B:ATP5C ratio to reflect the number of variants present in each pool. A ‘wildtype’ pool was generated by combing the parental plasmids in the same 5:8:8 ratio for use as a negative control.

### Mutant Library Transfection and Selection

HEK-293T landing pad cells ^99^ were maintained in glucose free DMEM (Gibco) supplemented with 10% heat inactivated FBS (Gibco), ±1% Pen/Strep, and 10 mM Galactose. On day −1, 500,000 cells with an ‘empty’ landing pad (LP-Neg) cells were plated into 6 well plates in antibiotic free media. On day 0, the cells were transfected with 1 μg of mutant library and 100 μg of *Bxb1* expressing plasmid (pCAG–NLS–HA–Bxb1; Addgene #51271, a gift from Pawel Pelczar) in triplicate. On day 1, the cells were replated into 60 mm TC dishes with antibiotic containing media and gene expression was induced with 1 μg/mL doxycycline. On day 3, cells were treated with 100 μg/mL Blasticidin S (Gibco) and 10 nM AP1903 (MedChemExpress) to select for cells which expressed the transgene and had undergone successful integration; selection was continued for 7 days. On day 10, HEK-293-integrated cells were plated into 6 well TC plates and treated with DMSO, apoptolidin A, ammocidin A at various concentrations (final DMSO concentration 0.25% v/v). Selective pressure was applied for two rounds of 4 days of exposure to each compound, followed by 3 days for recovery in glycomacrolide free media. After selection, surviving cells were allowed to expand until confluent at which point genomic DNA was extracted using the Qiagen DNeasy Blood & Tissue Kit.

### Amplicon Preparation and Sequencing

The integrated ATP synthase genes were PCR amplified using primers designed to anneal to the landing pad (TRE3G_fwd) and plasmid IRES sequence (IRES_rev) using 5% of the genomic DNA as a template and Q5 polymerase (New England Biolabs - NEB) at 100 μL scale. The PCR product was purified using the Monarch PCR & DNA clean-up kit (NEB). All library preparation and sequencing was carried out at the Vanderbilt Technologies for Advanced Genomics (VANTAGE) center. Sequencing libraries were prepared by tagmentation using the Nextera Flex library preparation kit (Illumina Inc. San Diego, CA, USA) using the manufacturers protocol. PE150 sequencing was carried out on the Novaseq 6000 platform at VATNAGE. Data are deposited in the gene expression omnibus (GEO), GSM5224467.

### Analysis of Saturation Mutagenesis Experiments

Demultiplexed reads were down-sampled to 5M paired reads using seqtk, aligned to the reference ORFs using bwa-mem ^100^ with a gap opening penalty of 100, and converted to BAM format using samtools. Variant frequency was determined using gatk version 4.1.8.1 ^101^. AnalyzeSaturationMutagenesis was used for each ORF. The resulting amino acid frequency tables were analyzed using a custom R script with the tidy data deposited on GEO as a supplementary file. The log10 fold change for each amino acid was calculated by subtracting the log10 transformed frequency of each treated replicate to its parental frequency.

### Generation and Validation of CRISPR lines

Isogenic mutants were generated using CRISPR-Cas9 gene editing as described in ^44^, using pSpCas9(BB)-2A-GFP (PX458), a gift from Feng Zhang (Addgene plasmid # 48138). Guide sequences (see supplement) were generated by annealing complementary oligonucleotides and golden-gate cloning with BbsI into PX458 and confirmed by sanger sequencing. 101–110 nt ssODN HDR templates were ordered with 50 bp of homology on either side of the cut site. HDR templates were designed to eliminate the PAM sequence to prevent re-cutting and designed to incorporate one or more synonymous mutations to add or remove restriction sites to facilitate screening of clones. 200,000 cells were electroporated using the Neon Transfection system (ThermoFisher), with 500 ng of PX458 plasmid ± 10 pmol ssODN template. Single cells were FACS sorted into U-bottom 96 well plates into conditioned media, selecting GFP+/PI-negative cells. Clones were expanded and genotyped by PCR of the targeted region, followed by restriction digest to identify edited clones. Successfully edited clones were further confirmed by PCR cloning and sanger sequencing of 6–8 colonies. ATPIF-1 K.O. lines were generated using PX458 without a donor template as described above. Individual clones were screened using intracellular flow cytometry against ATPIF1.

### *In vivo* Murine Modeling

All animal experiments were conducted in accordance to guidelines approved by the IACUC at Vanderbilt University Medical Center. Male NSGS [NOD-scid IL2Rgnull3Tg (hSCF/hGM-CSF/hIL3)] mice (The Jackson Laboratory), 6 - 8 weeks old were irradiated with 100 cGy microwave radiation. Twenty-four hours later, mice were transplanted with 1 × 10^6^ MV-4-11 cells via tail vein injections in each irradiated mouse. Mice were randomized post xenograft transplantation into cages of 5. Prior to treatment, peripheral microchimerism was documented at week 1. Upon establishing microchimerism, mice were treated with either 0.1 mg/kg or 0.03 mg/kg ammocidin in saline or saline vehicle i.p. for 5 days on, 2 days off for 2 weeks. Murine CBC was analyzed from blood collected into EDTA tubes (Greiner Bio-One) and analyzed with a Hemavet (Drew Scientific) analysis system.

### Flow Cytometry

For analysis of MV-4-11 cells expressing the Perceval HR reporter, cells were treated with compounds for 16 hrs in 96 well plates in 100 μL of media. Prior to analysis, propidium iodide was added at a final concentration of 300 ng/mL as a viability stain. Perceval-HR fluorescence was recorded on the YFP and AmCyan channels and intact, single, PI-negative cells were selected for analysis. ATP-ADP ratio was calculated by subtracting the asinh transformed YFP and AmCyan intensities. Two independent clones were analyzed as biological replicates. Cells were analyzed using a 5-laser LSR instrument (Becton Dickinson).

For analysis of pS6 suppression and ATPIF1 expression, cells were analyzed using intracellular flow cytometry multiplexed using fluorescent cell barcoding. For pS6 suppression, cells were treated for 16 hrs with the indicated compound in 96 well plates, ATPIF1 expression was measured in untreated cells. At the time of analysis, Alexa Fluor 700-NHS ester was added to each well as a viability stain (final 20 ng/mL) and allowed to incubate for 15 min, cells were fixed with 1.6% PFA for 10 min, and permeabilized with ice cold methanol for 30 min at −20 °C. Barcoding and staining was performed as described previously^102^ and demultiplexed using DebarcodeR^103^.

For analysis of mouse xenografts, red blood cells were lysed with EL Buffer on ice (Qiagen), with remaining cells washed and resuspended in 1x PBS with 1% BSA and stained for 15 minutes with the following antibodies: human CD45-APC (Clone 2D1) (Biolegend), human CD33-PE-Cy7 (Clone P67.6) (Biolegend), murine CD45-PE (Clone 30-F11) (Biolegend) and DAPI (Biolegend). Cells were washed and submitted for flow cytometric analysis using a 3-laser LSRII (Becton Dickinson).

### Histology and Immunohistochemistry

Tissues were fixed in 4% paraformaldehyde for 48 hours and stored in 70% ethanol before being embedding in paraffin and sectioned at 5 μm. The bone tissue was decalcified prior to being embedded in paraffin. Sections were de-waxed in xylene and rehydrated in successive ethanol baths. Standard Mayer’s Hematoxylin and Eosin (H&E) staining was performed. Antigen retrieval using a standard pH 6 sodium citrate buffer (BioGenex) was performed and sections were stained with Monoclonal Mouse Anti-Human CD45 (Dako, M0701, dilution 1:200) using the M.O.M. Kit (Vector).

### Pharmacokinetic Studies

All animal experiments were conducted in accordance with guidelines approved by the IACUC at Vanderbilt University Medical Center. Pharmacokinetics of Ammocidin in NSGS male mice in biological triplicate were assessed in whole blood after dosing with ammocidin alone i.p. 0.5 mg/kg. Whole blood samples were collected up to in EDTA tubes for analysis of plasma. Blood plasma was mixed 1:1 with an internal standard solution consisting of 1 μM apoptolidin A in PBS. Metabolites were extracted with 200 μL of ethyl acetate, evaporated to dryness and resuspended in 50 μL of MeOH. Ammocidin A concentration was determined using LC-MS (Thermo TSQ Quantum Access Max) with technical duplicates on a 50 × 1.8mm C18 column, isocratic 60/40 H2O/Acetonitrile + 10 mM ammonium acetate at 250 μL/min in ESI+ mode monitoring ammocidin A (1139.7 → 208.8, CE 19 V, RT = 1.00 min) and apoptolidin A (1146.68 → 805.46, CE 19 V, RT = 1.66 min).

---

**Extended Data Fig. 1.**
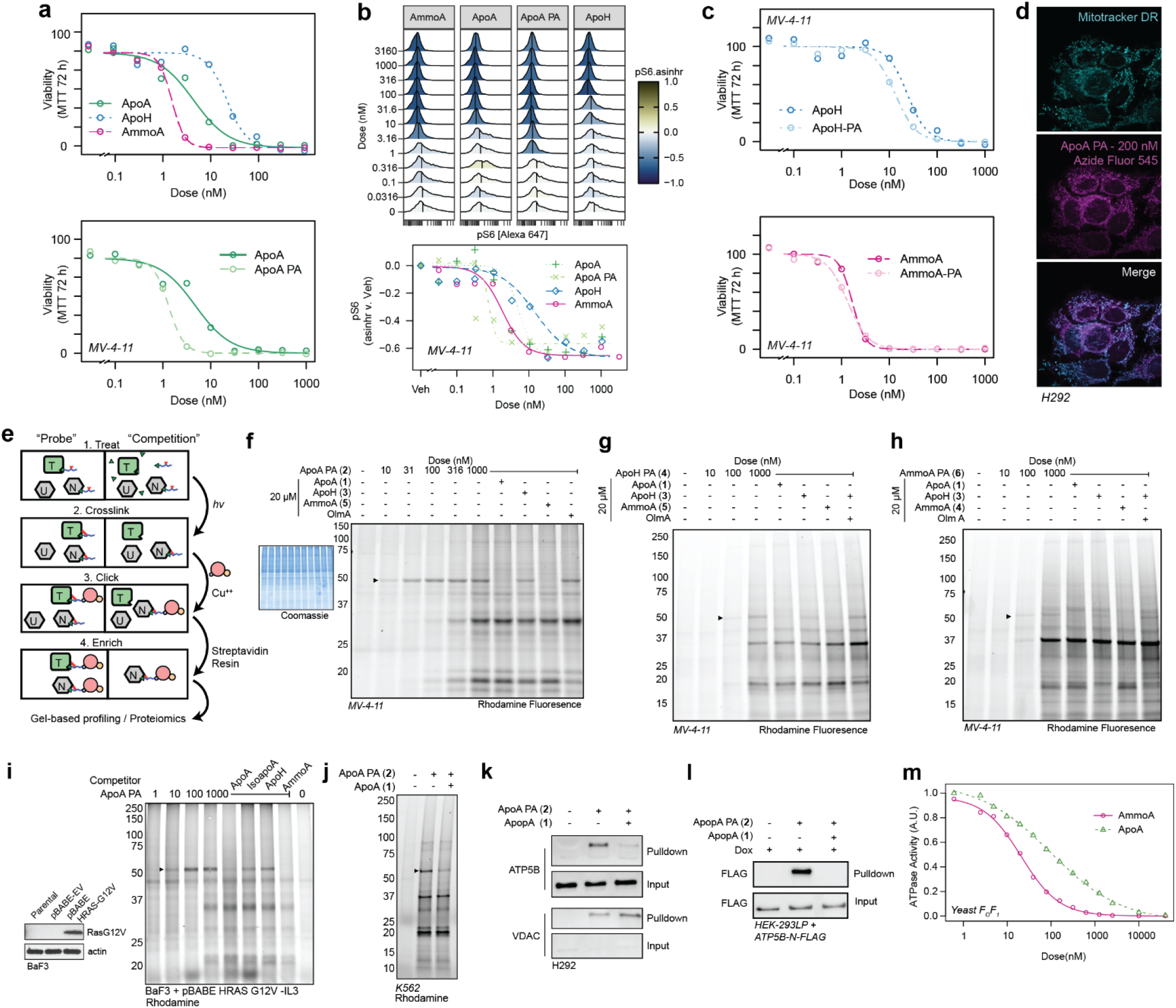
Validation of Glycomacrolide Photoaffinity Probes. **a,** Concentration response curves in MV-4-11 cells for apoptolidin family glycomacrolides (top) and probes compounds (bottom) at 72 h by MTT assay showing retention of activity; **b,** Analysis of pS6 phosphorylation using FCB barcoded flow cytometry of MV-4-11 cells treated for 16 h with glycomacrolides or probes; **c,** Concentrations response curves in MV-4-11 cells treated for 72 h with ApoH, AmmoA, or probes; **d,** Confocal microscopy of H292 cells treated with 200 nM ApoA PA for 1 h, photocrosslinked, fixed, and conjugated with rhodamine azide, showing mitochondrial localization of ApoA PA; **e,** Schematic of affinity enrichment workflow and competition experiments use to distinguish specific and non-specific binding partners – (T = [specific] target, N = non-specific binder, U = unbound protein); **f,** Gel-based profiling of ApoA PA adducts in MV-4-11 cells visualized using TAMRA-azide-desthiobiotin; **g,** Gel-based profiling of ApoH PA or **h**, AmmoA PA adducts in MV-4-11 cells; **i,** Gel-based profiling of ApoA PA targets in BaF3 cells transformed with HRas-G12V or **j**, K562 cells; **k,** Immunoblot showing specific enrichment of ATP5B in H292 cells treated with ApoA PA; **l,** Immunoblot showing pulldown of FLAG tagged ATP5B expressed in H292 landing pad cells (Fig. 3a); **m**, Concentration response-curves showing inhibition of ATPase activity in purified yeast F_O_F_1_ ATP synthase, measured using PK/LDH coupled assay.

**Extended Data Fig. 2.**
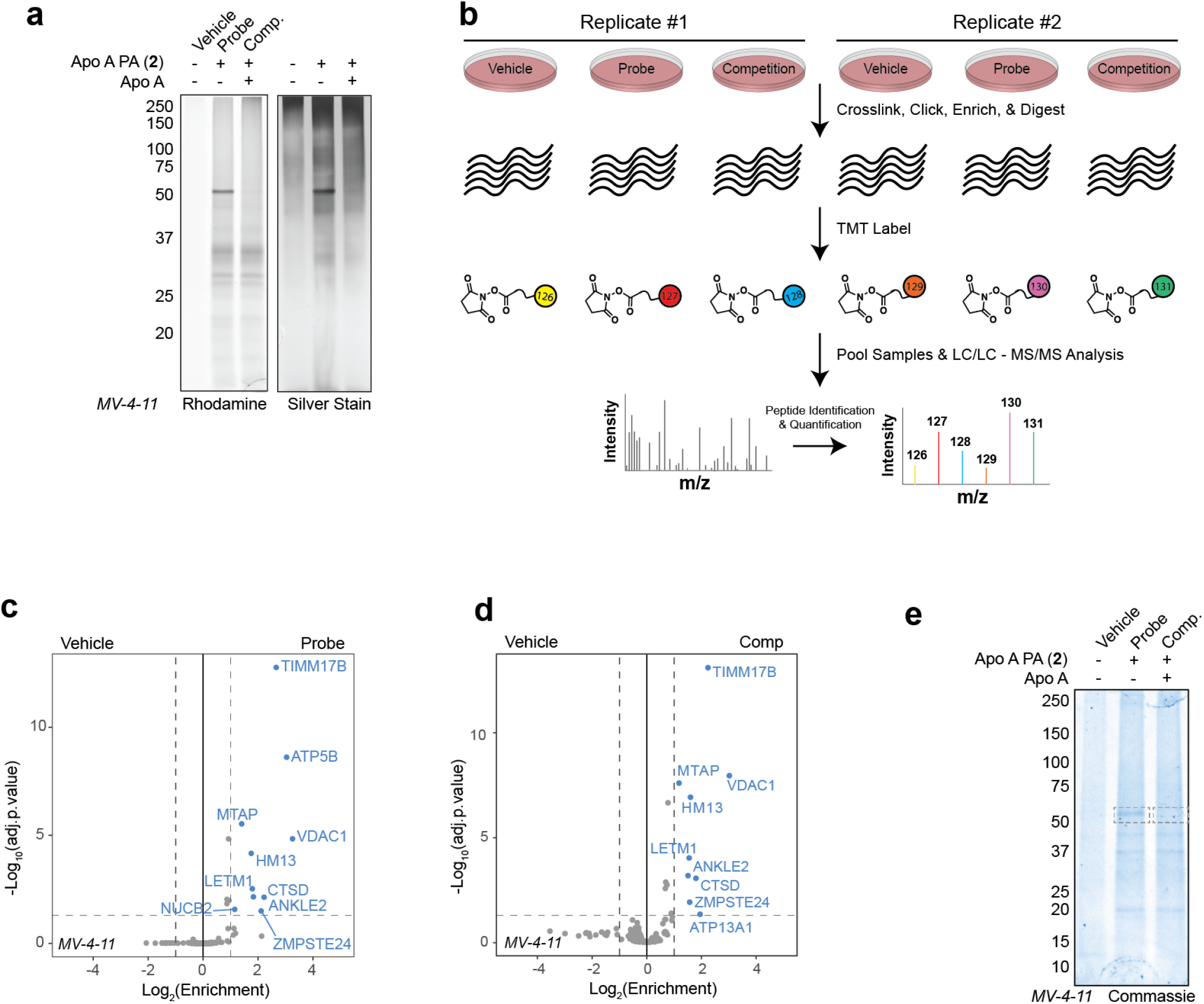
Identification of ATP5B as the target of apoptolidin A using comparative proteomics. **a,** Gel-based profiling of (**2**) targets in MV-4-11with ± 1 μM ApoA PA, ± 20 μM ApoA after affinity enrichment using streptavidin resin prior to digestion and TMT labeling imaged by rhodamine fluorescence and after silver staining; **b,** schematic of the TMT multiplexing strategy for to combine vehicle, probe, and competition, conditions from two separate biological replicates **c,** Volcano plot of proteins significantly enriched in ‘Probe’ condition compared to ‘Vehicle’ analyzed by TMT-multiplexed MuDPIT proteomics; **d,** Volcano plot of proteins significantly enriched in ‘Competition’ condition compared to ‘Vehicle’ analyzed by TMT-multiplexed MuDPIT proteomics; **e,** Coomassie stained gel-based profiling of (2) targets in MV-4-11 cells – 50 kDa band from probe and competition conditions were cut and subjected to in-gel digestion (see Data S1).

**Extended Data Fig. 3.**
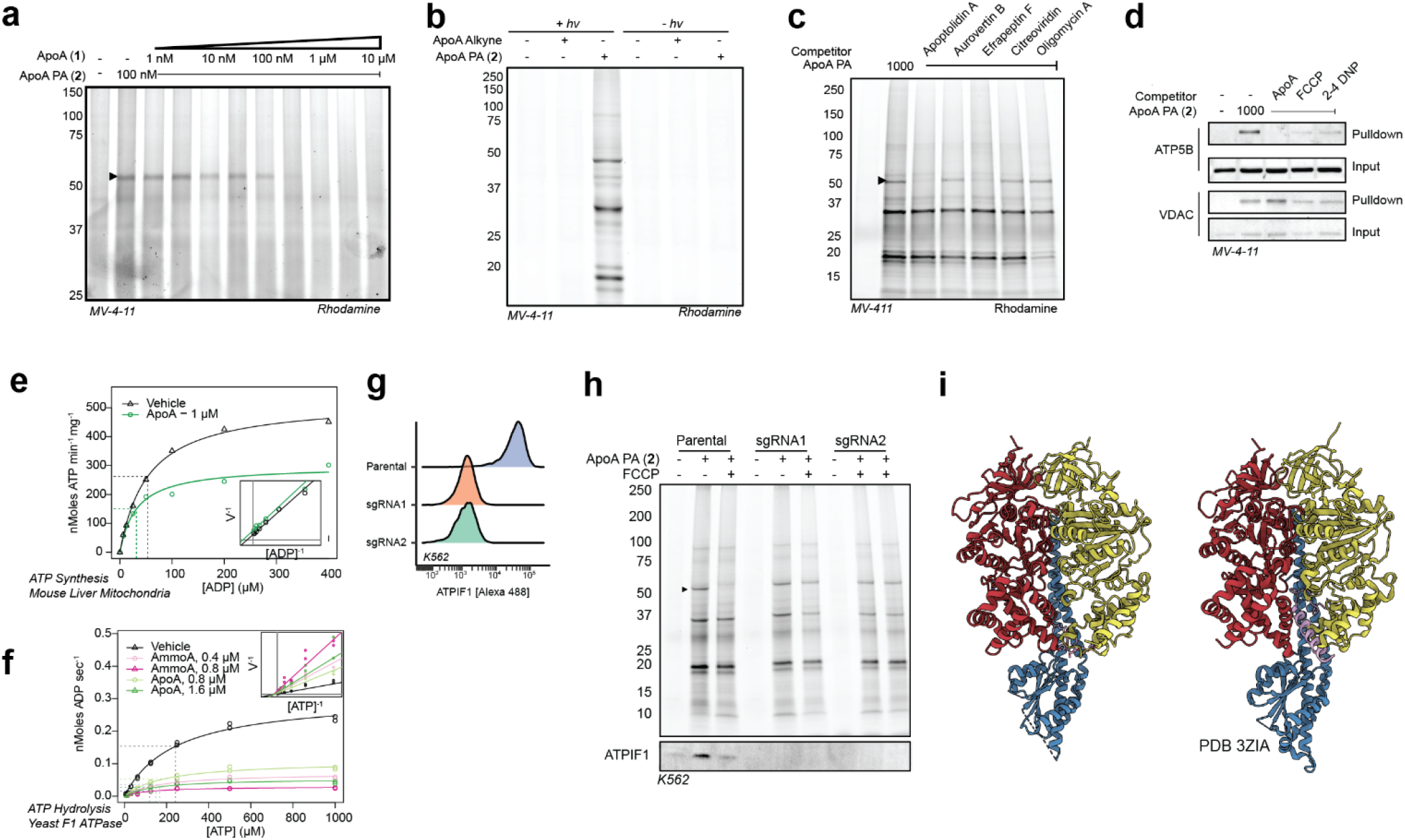
Analysis of apoptolidin A binding mode via photoaffinity labeling, and enzymatic assays. **a,** Gel-based analysis of apoptolidin A (**1**) competition against (**2**) at a fixed concentration of 100 nM (2); **b,** Gel-based analysis of diazirine and UV light dependent labeling of (2) at 1 μM; **c,** Gel-based analysis of (2) in the presence of known ATP synthase inhibitors; **d,** Gel-based profiling of ApoA PA in MV-4-11 cells treated with or without uncoupling agents (1 μM). **e,** Analysis of ADP concentration dependence on inhibition of ATP synthesis by apoptolidin A in isolated mouse liver mitochondria using HK/G6PDH coupled assay; **f,** Analysis of ATP concentration dependence on inhibition of ATP hydrolysis by (1) and (3) in isolated yeast F1 ATPase using a PK/LDH coupled assay. **g,** confirmation of ATPIF1 KO in two independent K562 clones after knockout using PX458 with two independent *ATP5IF1* targeting gRNAs; **h**, Gel-based profiling of membrane potential dependent adduction of ApoA PA in WT or IF1 KO K562 cells; **i,** comparison of ammocidin bound (left) to IF1 bound F_1_ ATPase.

**Extended Data Fig. 4.**
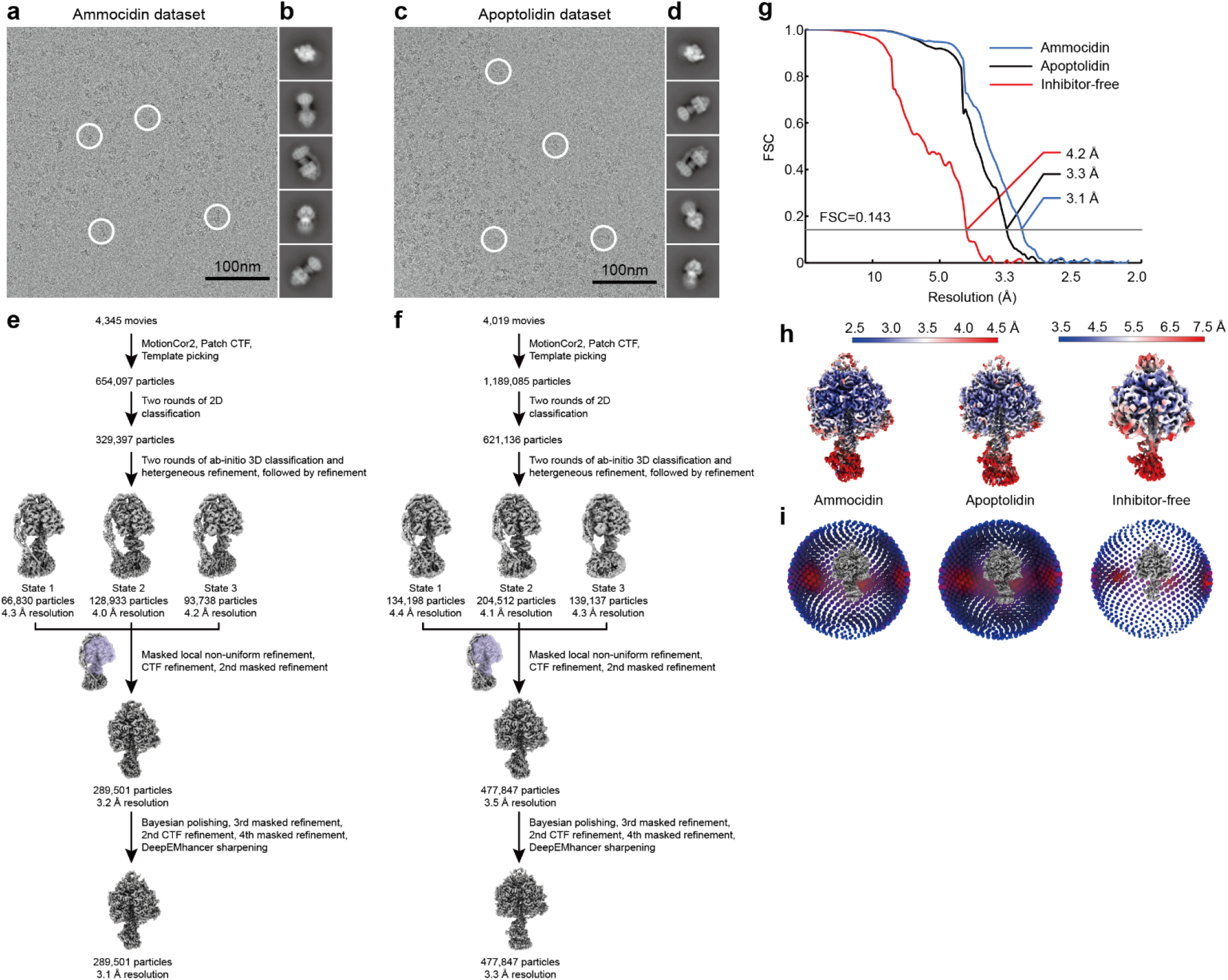
Workflow for cryoEM image analysis and validation for ammocidin-bound and apoptolidin-bound yeast ATP synthase. Example micrographs (**a**, **c**), 2D class average images (**b**, **d**), and workflow for obtaining maps of the F1 regions (**e**, **f**) for ammocidin-bound, and apoptolidin-bound ATP synthases, respectively. Corrected Fourier shell correlation curves after gold-standard refinement **a,** local resolution maps **b,** and orientation distribution plots **c,** are shown for the maps of the F1 regions of the ammocidin-bound, apoptolidin-bound, and inhibitor-free datasets.

**Extended Data Fig. 5.**
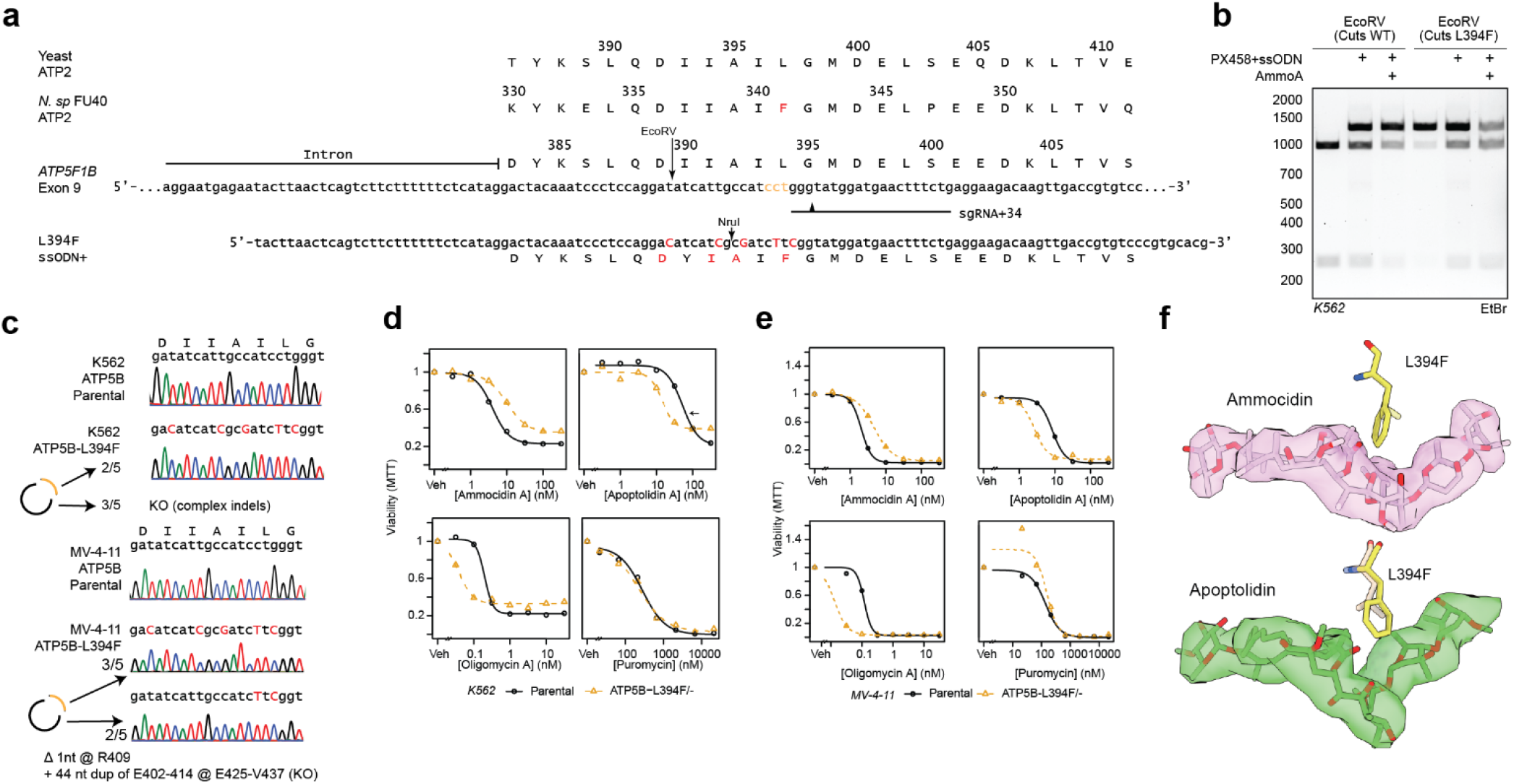
ATP5B-L394F mutation has opposing effects on apoptolidin and ammocidin sensitivity. **a,** Multiple sequence alignment of ATP synthase β subunit C-terminal domain and CRISPR/Cas9 HDR editing strategy for human exon 9 and using plasmid (PX458) encoding sgRNA and Cas9 and ssODN repair template, PAM sites labeled in yellow, cut locations noted with arrowhead, edits noted in red, restriction sites noted; **b**, Restriction digest of ATP5F1B Exon 9/10 amplicons from parental K562s or PX458 + ssODN treated K562s treated with ammocidin A (150 nM × 5 d) showing loss of wildtype allele and retention of edited alleles with ammocidin treatment; **c,** Confirmation of introduction of ATP5B-L394F mutation and KO of wildtype ATP5B in K562s and MV-4-11 cells by Sanger sequencing; **d,** ATP5B-L394F sensitizes cells to apoptolidin A and provides some resistance against ammocidin in K562 and **e,** MV-4-11 cells as measured by MTT assay at 72 hrs; **f,** Modeling of the L394 (yeast L397) mutation on the ammocidin (top) and apoptolidin (bottom) structures showing proximity to the hemiketal.

**Extended Data Fig. 6.**
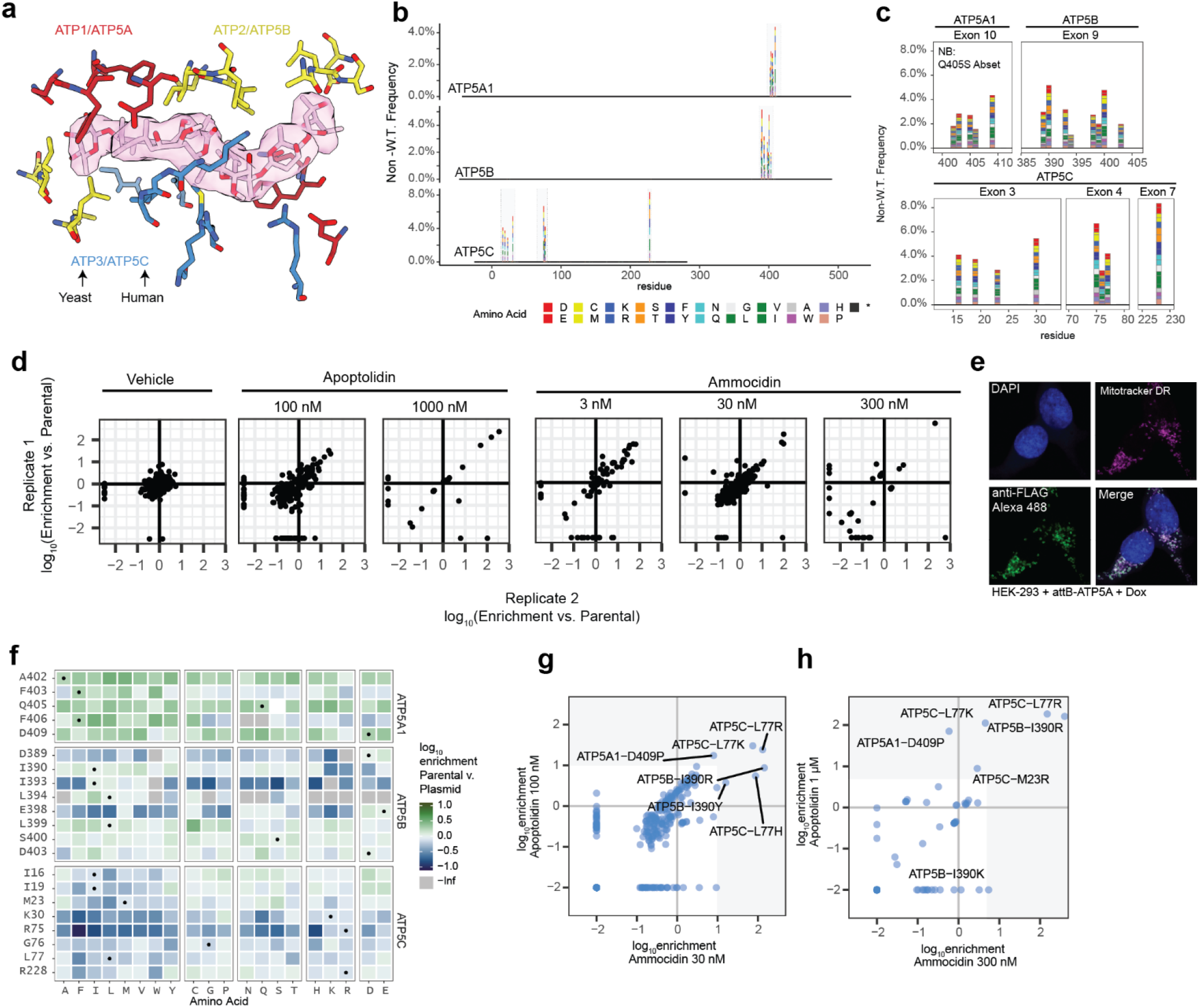
Deep Mutational Scanning of Ammocidin Binding Site on ATP Synthase. **a,** Selection of residues for mutagenesis on α (Yeast ATP1, Human ATP5A – red), β (Yeast ATP2, Human ATP5B – yellow), and γ (Yeast ATP3, Human ATP5C – blue) subunits within 4.5 Å of ammocidin or apoptolidin; **b, c,** Validation of ATP synthase variant library after nicking mutagenesis showing the frequency of non-wildtype variants at each position, with residue 1 corresponding to the first residue of the mature peptide; **d,** Assessment of reproducibility between biological replicates showing consistent enrichment of resistance mutations, variants which were not observed were set to −2.5; **e,** Confirmation of proper mitochondrial localization of the landing-pad expressed ATP5A by immunofluorescence at 63x in HEK-293T landing pad cells; **f,** Comparison of variant frequencies observed after selection for successful integration compared to the original library, variants which were observed in the original library but not detected after integration were set to -Inf. Variants which exhibit negative enrichment or complete loss are thought to exhibit decreased fitness relative to the WT alleles; **g,** Comparison of variants enriched by apoptolidin and ammocidin at high (1 μM / 300 nM) or **h,** medium dose (100 nM / 30 nM) demonstrating that apoptolidin and ammocidin select for similar resistance mutations.

**Extended Data Fig. 7.**
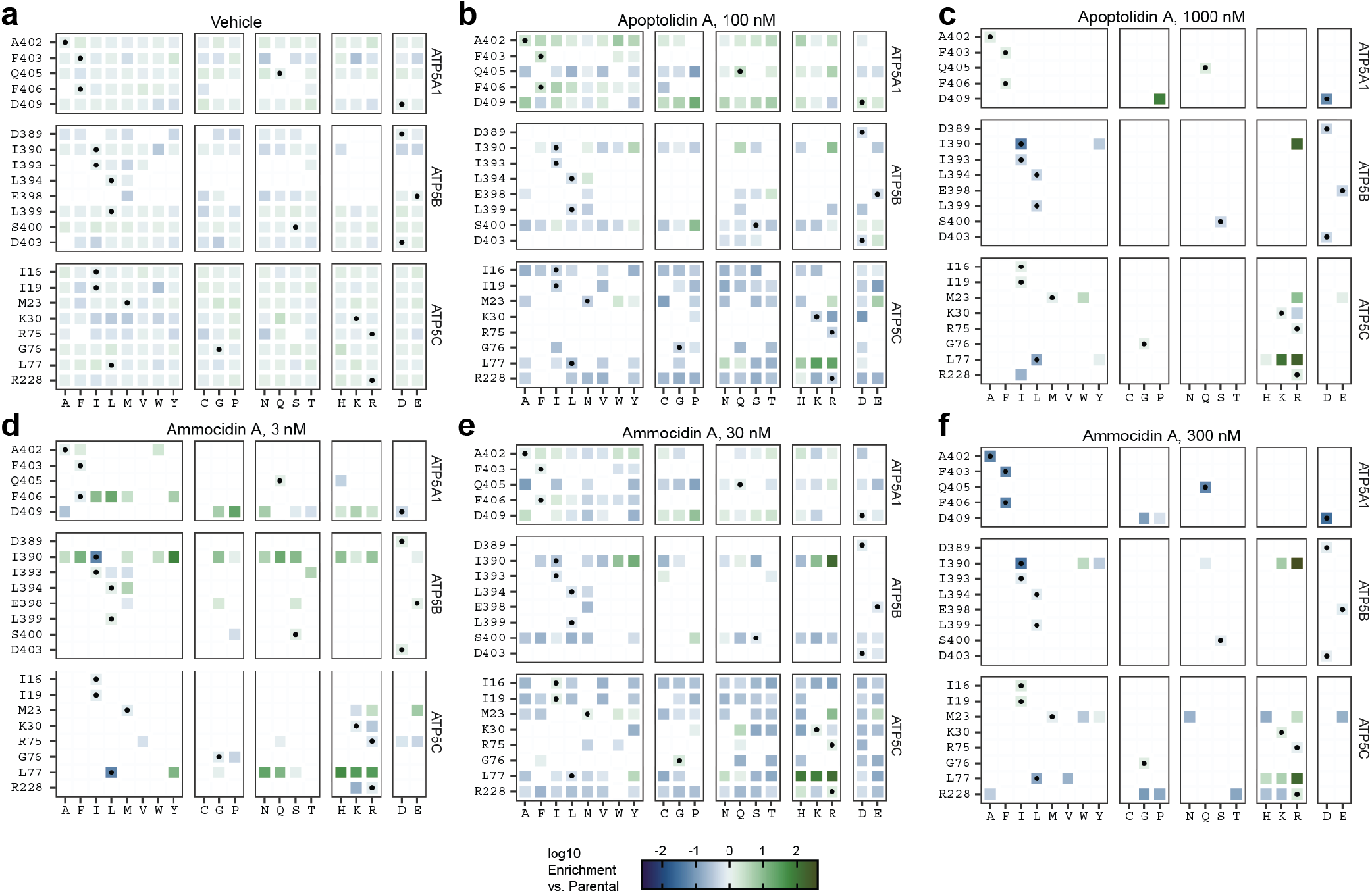
Apoptolidin and Ammocidin select for similar resistance mutations. Tile maps of amino acid enrichment relative to the parental library at each position at each dose of ammocidin or apoptolidin. Vehicle (**a**) treated cells were allowed to grow for one additional passage after selection. White squares indicate variants which were not detected (<100 counts) by deep sequencing. Circles represent WT alleles.

**Extended Data Fig. 8.**
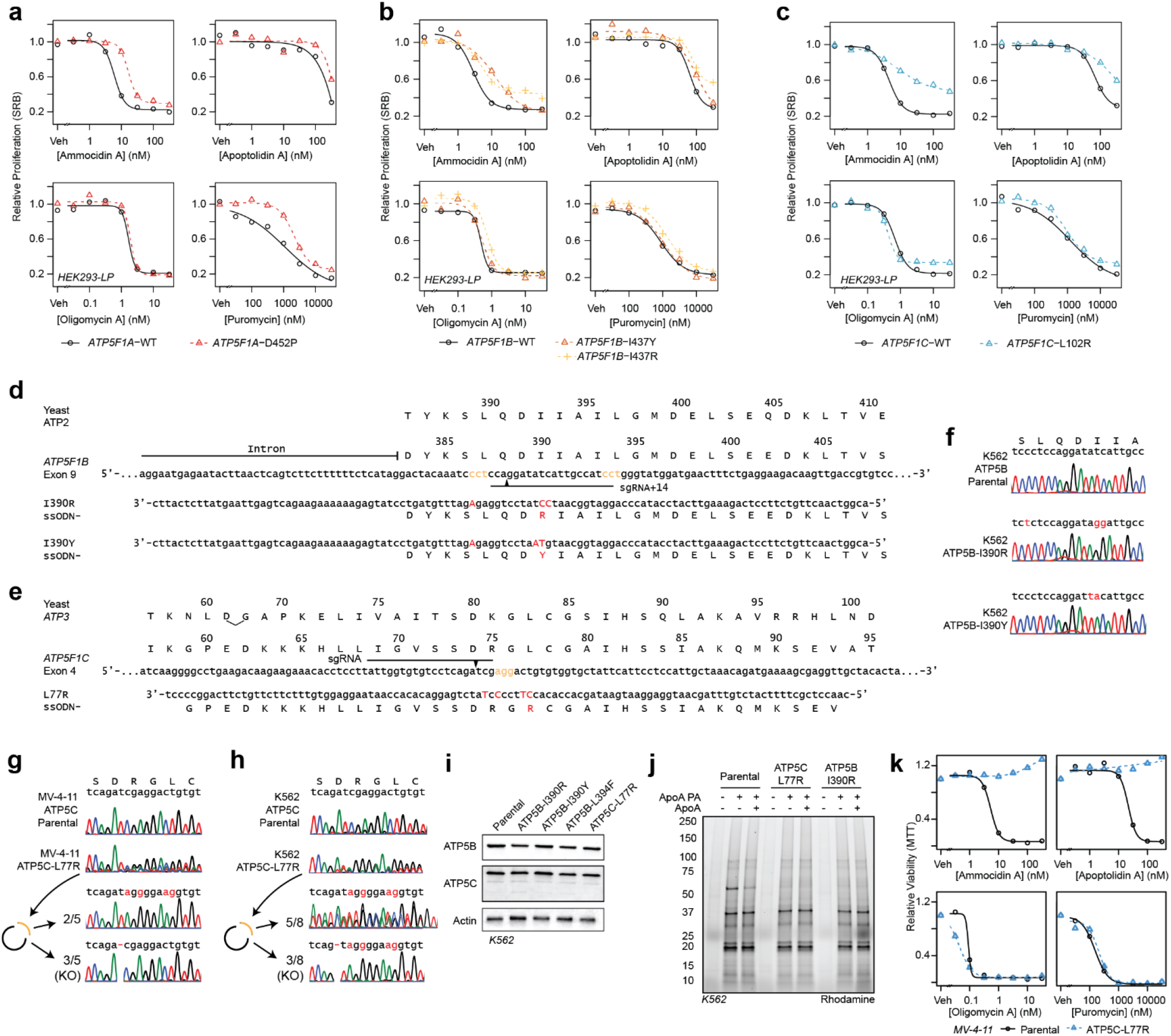
Validation of ATP synthase resistance mutations using CRISPR/Cas9 Genome Editing. **a,** Validation of resistance mutations using transgenic mutant ATP synthase in HEK293LP cells at 72 hrs by SRB assay for mutations in the alpha; **b,** beta; and **c,** gamma subunits; **d,** CRISPR/Cas9 HDR editing strategy for ATP5B exon 9 and **e,** ATP5C exon 4 using plasmid (PX458) encoding sgRNA and Cas9 and ssODN repair template, PAM sites labeled in yellow, cut locations noted with arrowhead, edits noted in red; **f,** Confirmation of introduction of ATP5B-I390R in K562 cells, **g,** ATP5C-L77R in MV-4-11 cells, or **h,** ATP5C-L77R in K562 cells by sanger sequencing of single-cell clones via PCR amplicons and/or PCR cloning to resolve heterozygous edits; **i,** Confirmation of unchanged expression of ATP synthase genes in CRISPR/Cas9 edited clones by immunoblot; **j**, Gel-based profiling of (2) targets in K562s Parental or CRISPR/Cas9 edited cell lines showing loss of binding to ATP5B; **i,** Secondary validation of ATP5C-L77R mutant in CRISPR edited MV-4-11 cell line using MTT-assay at 48 hrs.

**Extended Data Fig. 9.**
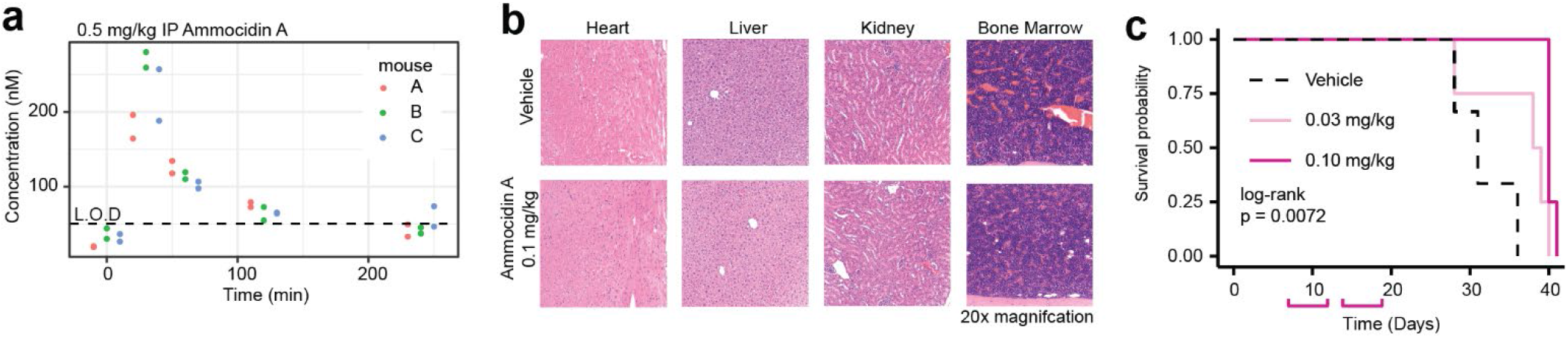
Evaluation of Ammocidin Dosing, Pharmacokinetics and Toxicity in NSGS Mice and efficacy in NSGS MV-4-11 Xenograft. **a,** Preliminary (non-GLP) evaluation of PK profile of ammocidin A in NSGS mice with a single dose of Ammocidin A (technical duplicate); **b,** Gross histology of H&E sections from organs of NSGS mice 2 weeks after treatment with ammocidin A (0.1 mg/kg IP QD for 5 on, 2 off) revealing/reveal no tissue damage related toxicity **c,** Engrafted mice (n = 4 per group) were treated from day 7 to day 18 (pink brackets) and therapy was then held. Mouse survival was measured by Kaplan-Meier analysis (p-value calculated using log-rank test.)

