## Supplementary Information for "A Family of Glycosylated Macrolides Selectively Target Energetic Vulnerabilities in Leukemia"

### Isolation of Apoptolidin A, Apoptolidin H, Ammocidin A.

Similar methods were used to obtain all three compounds by using different producing strains. Apoptolidin A was obtained by cultivation of wild-type *Nocardiopsis sp.* FU-40. Apoptolidin H was obtained by cultivation of *Nocardiopsis sp.* FU-40 in which *ApoGT2* was replaced with an apramycin resistance cassette as previously described[1]. Ammocidin A was obtained by cultivation of *Saccharothrix sp.* AJ9571 provided by Ajinomoto Co., Inc (Kawasaki, Japan). Each organism was plated on Bennett's agar (0.1% yeast extract, 0.1% beef extract, 0.2% N-Z Amine Type A, 1.0% dextrose, 2.0% agar, pH 7.0) and incubated at 30 °C for 3 - 7 days until sporulation. FU-40 Δ*ApoGT2* was grown on plates containing Bennet's media with the addition of apramycin, 80 µg/mL. The seed culture was initiated using spores scraped from the solid culture into 250 mL Erlenmeyer flasks containing 50 mL of seed medium (1.0% soluble starch, 1.0% molasses (Plantation Blackstrap, Unsulfured), 1.0% peptone, 1.0% beef extract, pH 7.0), and incubated for 7 days at 30 °C while shaking at 220 RPM. Production cultures were carried out in multiple 250 mL Erlenmeyer flasks containing 50 mL of production media (2.0% glycerol, 1.0% molasses, 0.5% casamino acids, 0.1% peptone, 0.4% calcium carbonate, pH 7.2) and incubated at 30 °C for 7 days while shaking at 220 RPM.

Apoptolidin A, apoptolidin H, and ammocidin A were purified using similar methods and differed only in the specific fractions collected according to their retention times. After 7 days of fermentation, the mycelia were separated from the culture broth by centrifugation at 3000 g x 30 min. The culture broth was extracted 3x with 1 volume of ethyl acetate and the combined organic layers were washed with brine, dried with $Na_2SO_4$ and concentrated *in vacuo*. The crude extracts were then subjected to chromatography with LH-20 resin using methanol as the mobile phase and the glycomacrolide containing fractions were identified by thin-layer-chromatography and pooled. The LH-20 fractions were then subjected to reverse phase HPLC using a Waters XBridge Prep C18 19 x 150 mm column with a 20-minute gradient from 70% A / 30% B to 20% A / 80% B, (Buffer A: 95% water, 5% acetonitrile, 10 mM ammonium acetate; Buffer B: 5% water, 95% acetonitrile, 10 mM ammonium acetate). Apoptolidin A – RT: 9.5 min, Apoptolidin H – RT: 8.0 min, Ammocidin A – RT 9.0 min. Fractions containing pure compounds were lyophilized using a Genevac HT-6 to yield white solids. Purity of each compound was confirmed by NMR ($^1$H and HSQC) and HPLC/MS.

### Isolation of Aurovertins B, D.

Aurovertin was obtained by cultivation of *Calcarisporium arbuscula,* NRRL-3705, obtained from the ARS Culture Collection, Peoria, IL, using the approach described by Baldwin [2]. Spores of NRRL-3705 were plated on modified Czapek agar (3.0% sucrose, 0.5% corn steep solids, 0.1% $KH_2PO_4$, 0.05% $MgSO_4$, 0.05% KCl, 0.01% $FeSO_4$) and incubated at 24 °C for 10 days. The production cultures were inoculated from a single colony of fungal mycelia into Roux bottles containing 70 mL of Baldwin production medium (3.0% Dextrose, 1.0% peptone, 0.5% NaCl, 0.5% dried corn steep, 0.5% yeast extract, 0.3% beef extract, pH 6.0) and incubated at room temperature for 14 days in the dark.

The fungal mycelia were separated from the broth by centrifugation and extracted with 250 mL of acetone for 2 hrs. The acetone was concentrated *in vacuo,* until only water remained. The aqueous layer was extracted 3 x with $CHCl_3$, the organic layer was washed with brine, and dried with $Na_2SO_4$ and concentrated to ~5 mL. The desired product was precipitated by the addition of pentane, filtered over Celite, and redissolved in methanol. The resulting semi-purified extract was then subjected to reverse phase HPLC using a Waters XBridge Prep C18 19 x 150 mm column with the following gradient at 10 mL /min : 0 min – 75 % A / 25 % B; 20 min – 60 % A/40 % B; 25 min – 60 % A/40 % B , 27 min – 5 %A/95 % B, 39 min – 5 % A/95 % B (Buffer A: 95 % water, 5 % acetonitrile, 10 mM ammonium acetate; Buffer B: 5% water, 95% acetonitrile, 10 mM ammonium acetate). Aurovertin D – RT: 15 min; Aurovertin B – RT: 30 min. Fractions containing pure compounds were lyophilized using a Genevac HT-6 to yield yellow solids. Purity of each compound was confirmed by NMR ($H^1$ and HSQC) and HPLC/MS.

**Isolation of Efrapeptins**

Efrapeptins were obtained by cultivation of *Tolypocladium cyclindrosporium,* ARSEF 962 (ATCC 42438) obtained from the American Type Tissue Culture Collection (ATCC, Manassas, VA). ARSEF 962 was plated on Sabouraud agar + yeast extract (4.0 % peptone, 1.0 % peptone, 1.0 %, 2.0 % agar, 1.0 % yeast extract, pH 5.6) and incubated at room temperature for 14 days. Production cultures were inoculated into 250 mL flasks containing 100 mL of Czapek Dox + peptone media (3.0% sucrose, 0.3 % $NaNO_3$, 0.1 % $KH_2PO_4$, 0.05 % $MgSO_4$, 0.05 % KCl, 0.01 % $FeSO_4$, 0.5 % peptone, pH 5.6) [3]. The fungal mycelia were separated from the broth by centrifugation and discarded. The broth was extracted 3x with dichloromethane, washed with brine, and dried with $Na_2SO_4$. The crude extract was concentrated *in vacuuo,* resuspended in 1 mL of methanol, and subjected to size exclusion chromatography using methanol as the mobile phase. The efrapeptin containing fractions were combined and subjected to reverse phase HPLC using a Waters XBridge Prep C18 19 x 150 mm column with the following gradient at 10 mL /min: 0 min – 70 % A / 30 % B; 20 min – 20 % A/80 %B (Buffer A: 95% water, 5 % acetonitrile, 10 mM ammonium acetate; Buffer B: 5% water, 95% acetonitrile, 10 mM ammonium acetate). Efrapeptin E – RT 16.8 min; efrapeptin F – RT 17.7 min. Fractions containing pure compounds were lyophilized using a Genevac HT-6 to yield white solids. Purity of each compound was confirmed by NMR ($H^1$ and HSQC) and HPLC/MS.

**Synthesis of Apoptolidin A PA (2)**

Photoaffinity probes were synthesized using reaction conditions adapted from Deguire, 2015 [4]. To a solution of 3-(3-(but-3-yn-1-yl)-3H-diazirin-3-yl)propanoic acid (4.5 mg, 0.027 mmol, Enamine, Kyiv, Ukraine) in dichloromethane (4.0 mL) on ice, was added bromo-tris-pyrrolidinophosphonium hexafluorophosphate (PyBrop, 13.3 mg, 0.28 mmol) and diisopropylethyl amine (DIPEA, 31 µL, 0.177 mmol). The resulting solution was stirred at 0 °C for 10 min. Apoptolidin A (20 mg 0.018 mmol) was added, followed by a crystal of 4-dimethylaminopyridine (DMAP). The resulting solution was warmed to room temperature overnight (16 h). The reaction was monitored by TLC (90:10 $CHCl_3$:MeOH) and quenched with 100 µL of MeOH and then concentrated. The resulting residue was diluted in EtOAc (20 mL) and washed with 1 M HCL (5 mL). The aqueous layer was extracted twice with EtOAC (2 x 10 mL). The organic extracts were combined and washed with $NaHCO_3$ (5 mL) and brine (5 mL), dried with anhydrous sodium sulfate and concentrated *in vacuo*. The resulting residue was dissolved in 800 µL of MeOH and purified by reversed phase HPLC using a Waters XBridge Prep C18 19 x 150 mm column, with a 15-minute gradient from 32 % to 85 % acetonitrile in water, with 25 mM ammonium bicarbonate. The fractions containing the desired product (rt = 11 min) were combined and lyophilized to afford (**2**) as a white solid (1.9 mg, 8% isolated yield). HRMS (ESI-TOF MS) m/z 1299.6973 $(M+Na)^+$ calculated, 1299.7023 observed (3.4 ppm). Regiochemistry of addition was confirmed by multidimensional NMR (see Table S1, Fig S13).

**Synthesis of Apoptolidin H PA (4)**

Apoptolidin H PA was synthesized using identical reaction conditions and work-up as used for (**2**), with Apoptolidin H (14.9 mg 0.018 mmol) The resulting residue was dissolved in 800 µL of MeOH and purified by reversed phase HPLC using a Waters XBridge Prep C18 19 x 150 mm column, with a 20-minute gradient from 32% to 77% acetonitrile in water, with 25 mM ammonium bicarbonate. The fractions containing the desired product (rt = 14 min) were combined and lyophilized to afford (**4**) as a white solid (2.4 mg, 13% isolated yield). HRMS (ESI-TOF MS) m/z 1011.5394 (M+Na)+ calculated, 1011.5405 observed (1.1 ppm). Regiochemistry of addition was confirmed by multidimensional NMR (see Table S2, Fig. S14).

**Synthesis of Ammocidin A PA (6)**

Ammocidin A PA was synthesized using identical reaction conditions and work-up as used for (**2**), with Ammocidin A (21 mg 0.018 mmol) The resulting residue was dissolved in 800 µL of MeOH and purified by reversed phase HPLC using a Waters XBridge Prep C18 19 x 150 mm column, with a 20-minute gradient from 32% to 77% acetonitrile in water, with 25 mM ammonium bicarbonate. The fractions containing the desired product (rt = 16 min) were combined and lyophilized to afford (**6**) as a white solid (0.6 mg, 2.5% isolated yield). HRMS (ESI-TOF MS) m/z 1287.6854 $(M+Na)^+$ calculated 1287.6866, observed (0.9 ppm). Regiochemistry of addition was confirmed by multidimensional NMR (see Table S3, Fig. S15).

### Table S1: NMR validation of Apoptolidin A PA

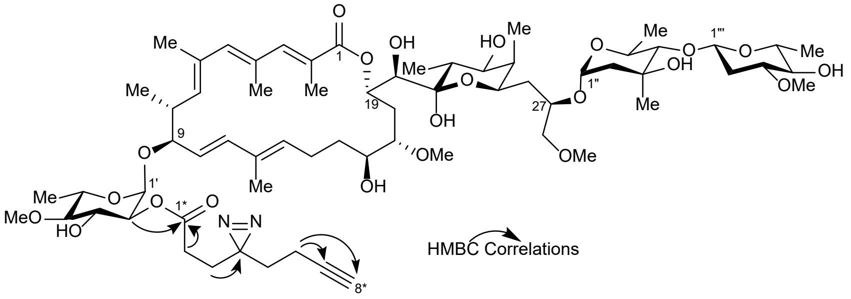

| | Apoptolidin A PA-Alkyne **(2)** | | | | |
|---|---|---|---|---|---|
| Position | $\delta_C$ | $\delta_H$ | Position | $\delta_C$ | $\delta_H$ |
| 1 | 172.3 | | 1' | 93.2 | 4.98 |
| 2 | 123.5 | | 2' | 75 | 4.56 |
| 3 | 149 | 7.38 | 3' | 72.1 | 3.9 |
| 4 | 132.9 | | 4' | 87.1 | 2.81 |
| 5 | 146.8 | 6.19 | 5' | 67.9 | 3.77 |
| 6 | 133.4 | | 6' | 18.1 | 1.28 |
| 7 | 142.5 | 5.21 | 4'-OMe | 61 | 3.59 |
| 8 | 38.9 | 2.68 | | | |
| 9 | 84.1 | 3.83 | 1" | 99.3 | 4.94 |
| 10 | 125.2 | 5.12 | 2" | 45.4 | 1.92, 1.8 |
| 11 | 140.9 | 6.14 | 3" | 72.7 | |
| 12 | 134.3 | | 4" | 85.6 | 3.35 |
| 13 | 133.5 | 5.63 | 5" | 67.1 | 3.67 |
| 14 | 24.3 | 2.45, 2.05 | 6" | 18.7 | 1.25 |
| 15 | 36.3 | 1.5, 1.42 | 3"-Me | 22.4 | 1.32 |
| 16 | 74.4 | 3.41 | | | |
| 17 | 83.5 | 2.76 | 1''' | 101.6 | 4.83 |
| 18 | 38 | 2.16, 1.74 | 2''' | 36.8 | 2.45, 1.27 |
| 19 | 72.2 | 5.3 | 3''' | 81.8 | 3.18 |
| 20 | 75.1 | 3.53 | 4''' | 76.9 | 2.97 |
| 21 | 101 | | 5''' | 73.1 | 3.22 |
| 22 | 36.1 | 2.05 | 6''' | 18.2 | 1.28 |
| 23 | 73.5 | 3.71 | 3'''-OMe | 57 | 3.42 |
| 24 | 40.5 | 1.73 | | | |
| 25 | 69.2 | 3.95 | 1* | 173.1 | |
| 26 | 36.7 | 1.58, 1.46 | 2* | 28.8 | 2.25, 2.16 |
| 27 | 76.6 | 3.44 | 3* | 28.7 | 1.82, 1.75 |
| 28 | 76.4 | 3.32 | 4* | 78.8 | |
| 2-Me | 13.9 | 2.1 | 5* | 33.2 | 1.62 |
| 4-Me | 17.7 | 2.19 | 6* | 13.6 | 2.04 |
| 6-Me | 16.3 | 1.94 | 7* | 83.4 | |
| 8-Me | 18.3 | 1.14 | 8* | 70.2 | 2.28 |
| 12-Me | 12 | 1.66 | | | |
| 22-Me | 11.9 | 1.03 | | | |
| 24-Me | 5 | 0.9 | | | |
| 17-OMe | 61 | 3.37 | | | |
| 28-OMe | 59.1 | 3.27 | | | |

Data recorded in $CD_3OD$ at 600 Hz

**Table S2: NMR validation of Apoptolidin H PA (4)**

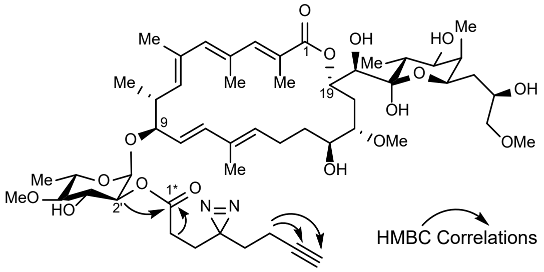

| | Apoptolidin H PA-Alkyne **(4)** | | | | |
|---|---|---|---|---|---|
| Position | $\delta_C$ | $\delta_H$ | Position | $\delta_C$ | $\delta_H$ |
| 1 | 172.6 | | 1' | 93.3 | 4.98 |
| 2 | 123.8 | | 2' | 75 | 4.55 |
| 3 | 148.7 | 7.36 | 3' | 72 | 3.91 |
| 4 | 133.5 | | 4' | 87 | 2.81 |
| 5 | 146.7 | 6.19 | 5' | 67.9 | 3.77 |
| 6 | 133.3 | | 6' | 17.7 | 1.29 |
| 7 | 142.5 | 5.22 | 4'-OMe | 60.9 | 3.59 |
| 8 | 38.7 | 2.76 | | | |
| 9 | 84 | 3.83 | | | |
| 10 | 126.1 | 5.22 | | | |
| 11 | 141 | 6.17 | | | |
| 12 | 134.8 | | | | |
| 13 | 133 | 5.68 | | | |
| 14 | 24.3 | 2.47, 2.06 | | | |
| 15 | 36.2 | 1.52, 1.41 | | | |
| 16 | 74.5 | 3.43 | | | |
| 17 | 83.5 | 2.73 | | | |
| 18 | 38.2 | 2.15, 1.75 | | | |
| 19 | 72 | 5.31 | | | |
| 20 | 75.1 | 3.54 | | | |
| 21 | 101 | | | | |
| 22 | 36.1 | 2.04 | | | |
| 23 | 73.5 | 3.75 | | | |
| 24 | 40.5 | 1.76 | | | |
| 25 | 68.9 | 4.09 | 1* | 173.2 | |
| 26 | 38.1 | 1.58, 1.29 | 2* | 28.8 | 2.26, 2.17 |
| 27 | 67.9 | 3.55 | 3* | 28.3 | 1.87, 1.75 |
| 28 | 78.5 | 3.19 | 4* | 73.1 | |
| 2-Me | 14 | 2.11 | 5* | 33.2 | 1.63 |
| 4-Me | 17.5 | 2.21 | 6* | 13.7 | 2.03 |
| 6-Me | 16.1 | 1.92 | 7* | 83.1 | |
| 8-Me | 18.1 | 1.14 | 8* | 70.3 | 2.28 |
| 12-Me | 11.8 | 1.67 | | | |
| 22-Me | 12.1 | 1.02 | | | |
| 24-Me | 5.1 | 0.89 | | | |
| 17-OMe | 61.4 | 3.36 | | | |
| 28-OMe | 59.3 | 3.29 | | | |

Data recorded in $CD_3OD$ at 600 Hz

### Table S3: NMR validation of Ammocidin A PA (6)

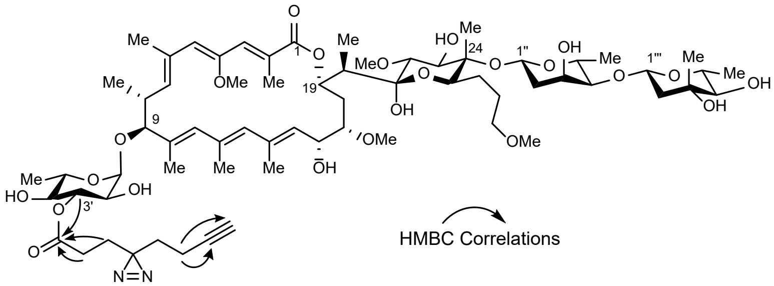

| | | | | | |
|---|---|---|---|---|---|
| | Ammocidin A PA-Alkyne **(6)** | | | | |
| Position | $\delta_C$ | $\delta_H$ | Position | $\delta_C$ | $\delta_H$ |
| 1 | 171.2 | | 1' | 95.6 | 4.7 |
| 2 | 125.4 | | 2' | 71.6 | 3.58 |
| 3 | 138.2 | 7 | 3' | 76.8 | 5.2 |
| 4 | 153.6 | | 4' | 87.1 | 3.15 |
| 5 | 134.2 | 5.79 | 5' | 68.7 | 3.86 |
| 6 | 131.8 | | 6' | 18.1 | 1.29 |
| 7 | 142 | 5.35 | | | |
| 8 | 36.8 | 2.88 | 1" | 92.6 | 5.11 |
| 9 | 87.9 | 3.85 | 2" | 38.9 | 1.97, 1.76 |
| 10 | 134.8 | | 3" | 68.3 | 4.28 |
| 11 | 136.3 | 5.94 | 4" | 83.3 | 3.31 |
| 12 | 134.1 | | 5" | 69.5 | 3.92 |
| 13 | 133.5 | 5.61 | 6" | 18.3 | 1.27 |
| 14 | 136.2 | | | | |
| 15 | 129.4 | 5.25 | 1''' | 101.8 | 4.72 |
| 16 | 67.9 | 4.83 | 2''' | 46.6 | 1.97, 1.74 |
| 17 | 82.8 | 2.7 | 3''' | 72.3 | |
| 18 | 35.8 | 1.93, 1.67 | 4''' | 80.2 | 3.13 |
| 19 | 72.3 | 5.54 | 5''' | 72.6 | 3.41 |
| 20 | 44.9 | 2.07 | 6''' | 19 | 1.3 |
| 21 | 99.9 | | 3'''-Me | 20.4 | 1.27 |
| 22 | 81.9 | 3.08 | | | |
| 23 | 76.8 | 3.95 | | | |
| 24 | 82.4 | | | | |
| 25 | 73.9 | 3.66 | 1* | 174 | |
| 26 | 25.8 | 1.60, 1.24 | 2* | 29.4 | 2.27 |
| 27 | 27.8 | 1.53, 1.34 | 3* | 29 | 1.78 |
| 28 | 73.9 | 3.3 | 4* | - | |
| 2-Me | 13.8 | 2.12 | 5* | 33.2 | 1.65 |
| 6-Me | 13.1 | 2.09 | 6* | 13.7 | 2.05 |
| 8-Me | 18.2 | 1.2 | 7* | 83.1 | |
| 10-Me | 12 | 1.62 | 8* | 70.7 | 2.27 |
| 12-Me | 18.1 | 1.91 | | | |
| 14-Me | 17.6 | 1.76 | | | |
| 20-Me | 9.3 | 1.14 | | | |
| 24-Me | 10.7 | 1.16 | | | |
| 4-OMe | 61.4 | 3.54 | | | |
| 17-OMe | 57.6 | 3.38 | | | |
| 22-OMe | 61.4 | 3.57 | | | |
| 28-OMe | 58.7 | 3.27 | | | |

Data recorded in $CD_3OD$ at 600 Hz

### Table S4: CryoEM data collection, refinement, and validation statistics

| | $yF_1F_O$ ammocidin (EMDB-23763) (PDB 7MD2) | $yF_1F_O$ apoptolidin (EMDB-23764) (PDB 7MD3) | $yF_1F_O$ (EMDB-23765) |
|---|---|---|---|
| **Data collection and processing** | | | |
| Magnification | 75,000 | 75,000 | 25,000 |
| Voltage (kV) | 300 | 300 | 200 |
| Electron exposure (e–/Å$^2$) | 43 | 45 | 36 |
| Defocus range (μm) | 0.7-2.4 | 1.1-2.5 | 1.0-3.2 |
| Movie pixel size (Å) | 1.03 | 1.03 | 1.45 |
| Final map pixel size (Å) | 1.03 | 1.2875 | 1.45 |
| Symmetry imposed | C1 | C1 | C1 |
| Initial particle images (no.) | 654,097 | 1,189,085 | 38,514 |
| Final particle images (no.) | 289,501 | 477,847 | 34,035 |
| Map resolution (Å) | 3.1 | 3.3 | 4.2 |
| FSC threshold | 0.143 | 0.143 | 0.143 |
| Map resolution range (Å) | 2.5-5.0 | 2.8-5.0 | 3.8-8.0 |
| **Refinement** | | | |
| Initial model used (PDB code) | 2XOK | 2XOK | |
| Model resolution (Å) | 3.1 | 3.5 | |
| FSC threshold | 0.5 | 0.5 | |
| Model resolution range (Å) | 330-3.1 | 330-3.5 | |
| Map sharpening *B* factor (Å$^2$) | Sharpened locally | Sharpened locally | |
| Model composition | | | |
| Non-hydrogen atoms | 23474 | 23031 | |
| Protein residues | 3115 | 3133 | |
| Ligands | 8 | 8 | |
| *B* factors (Å$^2$) | | | |
| Protein | 77.99 | 75.71 | |
| Ligand | 59.67 | 57.29 | |
| R.m.s. deviations | | | |
| Bond lengths (Å) | 0.008 | 0.006 | |
| Bond angles (°) | 0.732 | 0.674 | |
| Validation | | | |
| MolProbity score | 1.10 | 1.23 | |
| Clashscore | 3.07 | 3.65 | |
| Poor rotamers (%) | 0.25 | 0.26 | |
| Ramachandran plot | | | |
| Favored (%) | 97.99 | 97.65 | |
| Allowed (%) | 1.78 | 2.09 | |
| Disallowed (%) | 0.23 | 0.26 | |

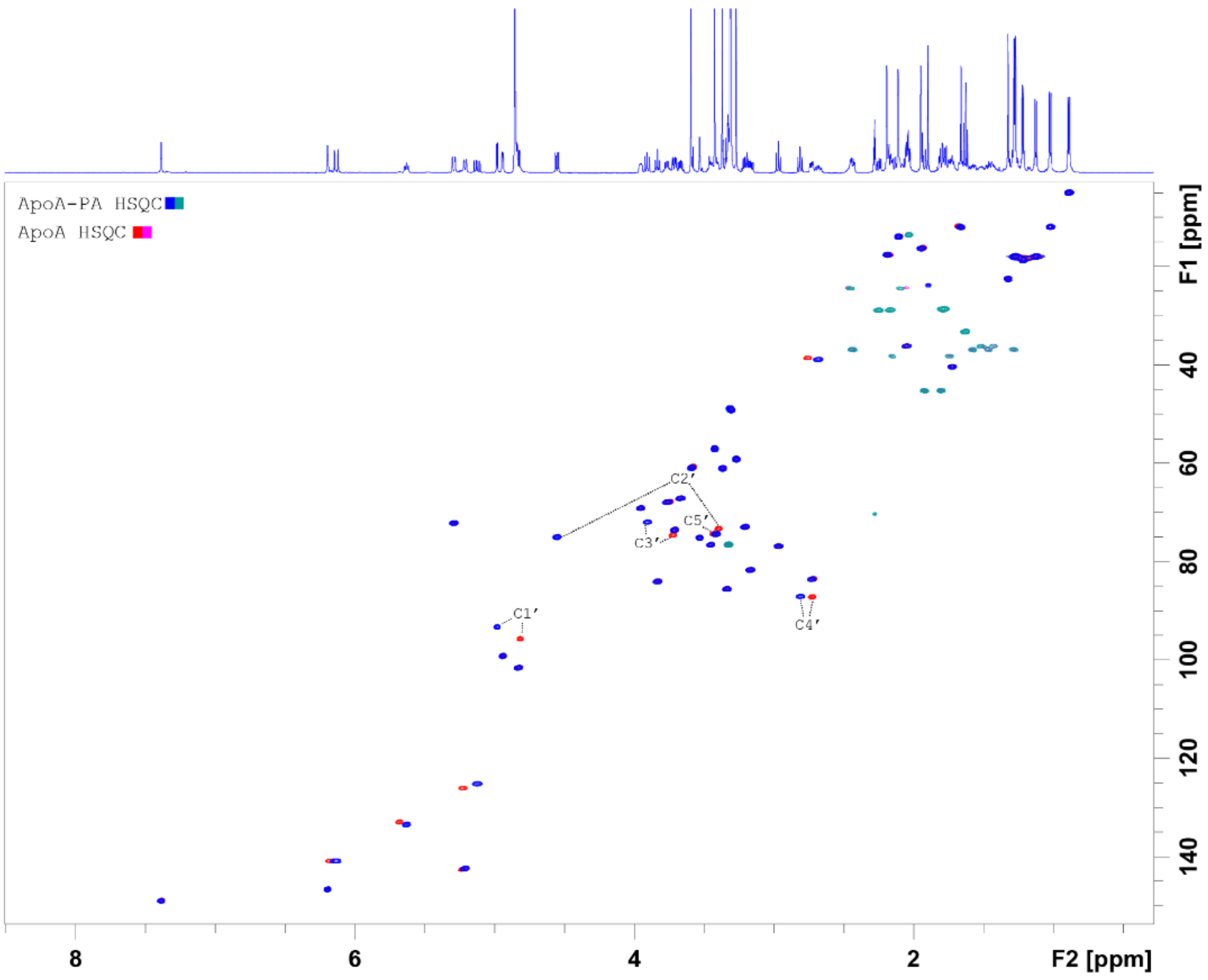

**Fig. S1 | HSQC of Apoptolidin A PA (2) in MeOD at 600 MHz**

Apoptolidin A PA (blue/green) on Apoptolidin A (red/pink). C9 6-deoxy-4-*O*-methyl-α-L-glucose carbons labeled

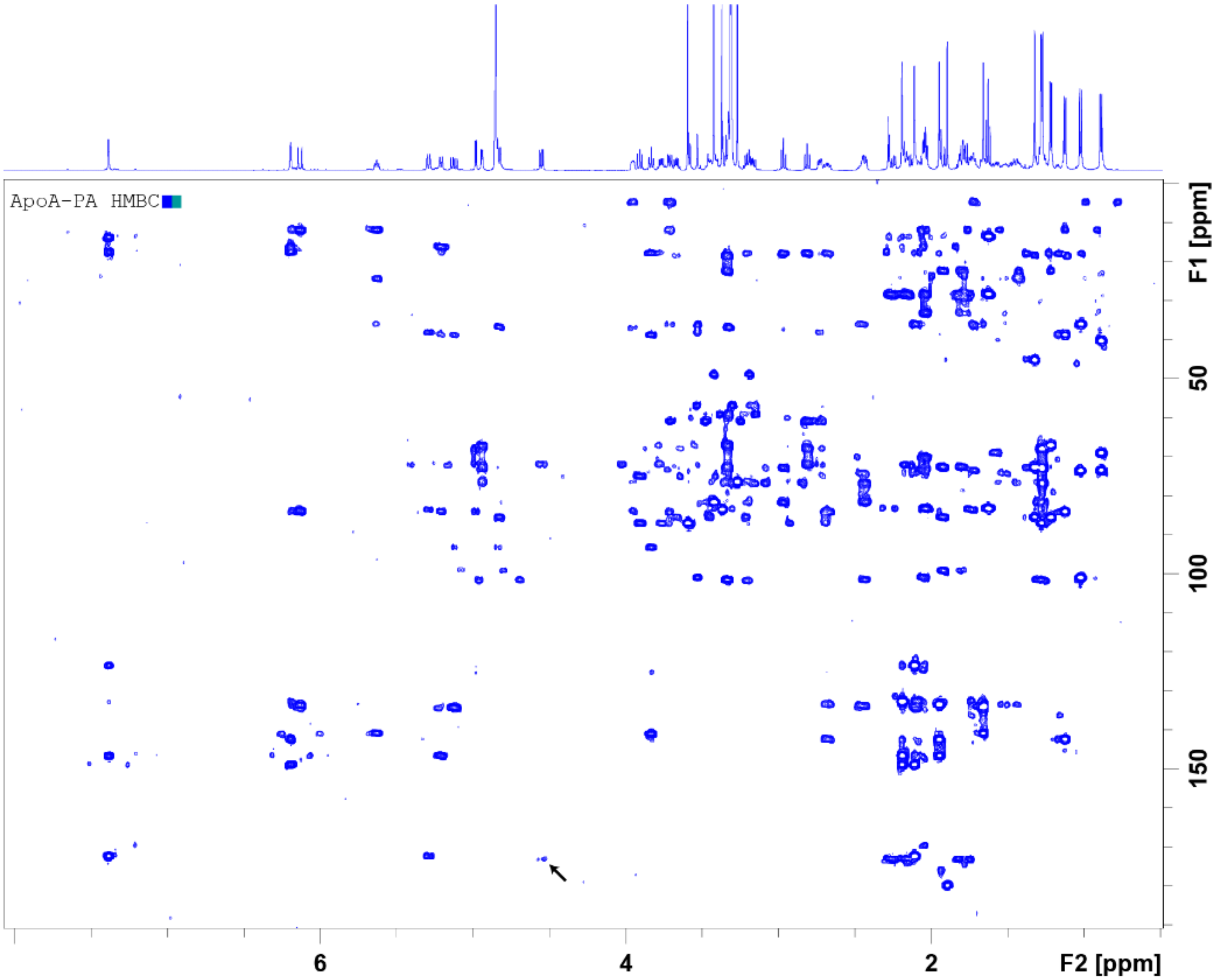

**Fig. S2 | HMBC of Apoptolidin A PA (2) in MeOD at 600 MHz**

HMBC spectrum of Apoptolidin A PA in MeOD at 600 MHz with correlation between C2’ and 1* (carbonyl of diazirine/alkyne ester) denoted by the arrow.

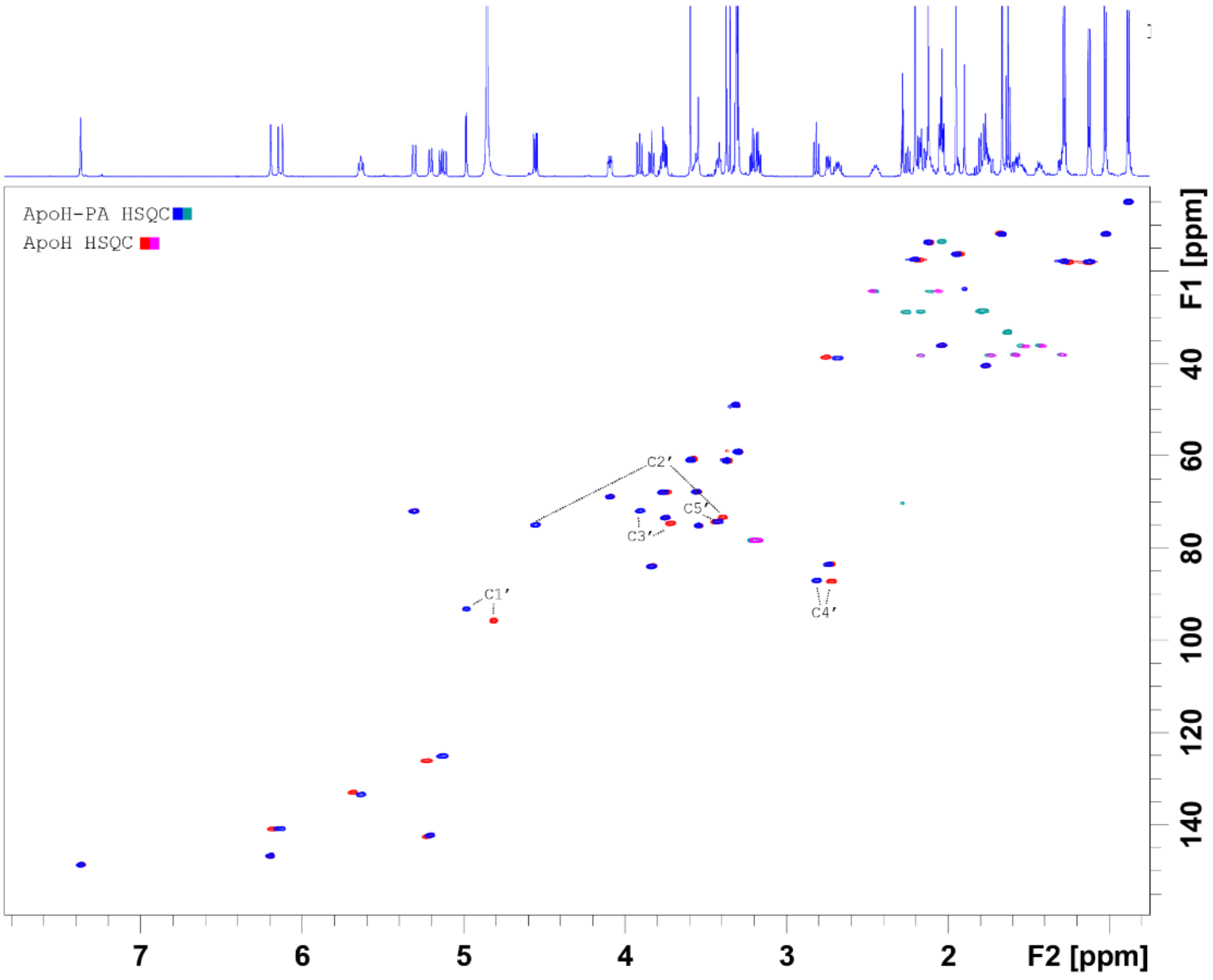

**Fig. S3 | HSQC of Apoptolidin H PA (2) in MeOD at 600 MHz**

Apoptolidin H PA (blue/green) on Apoptolidin H (red/pink). C9 6-deoxy-4-*O*-methyl-α-L-glucose carbons labeled

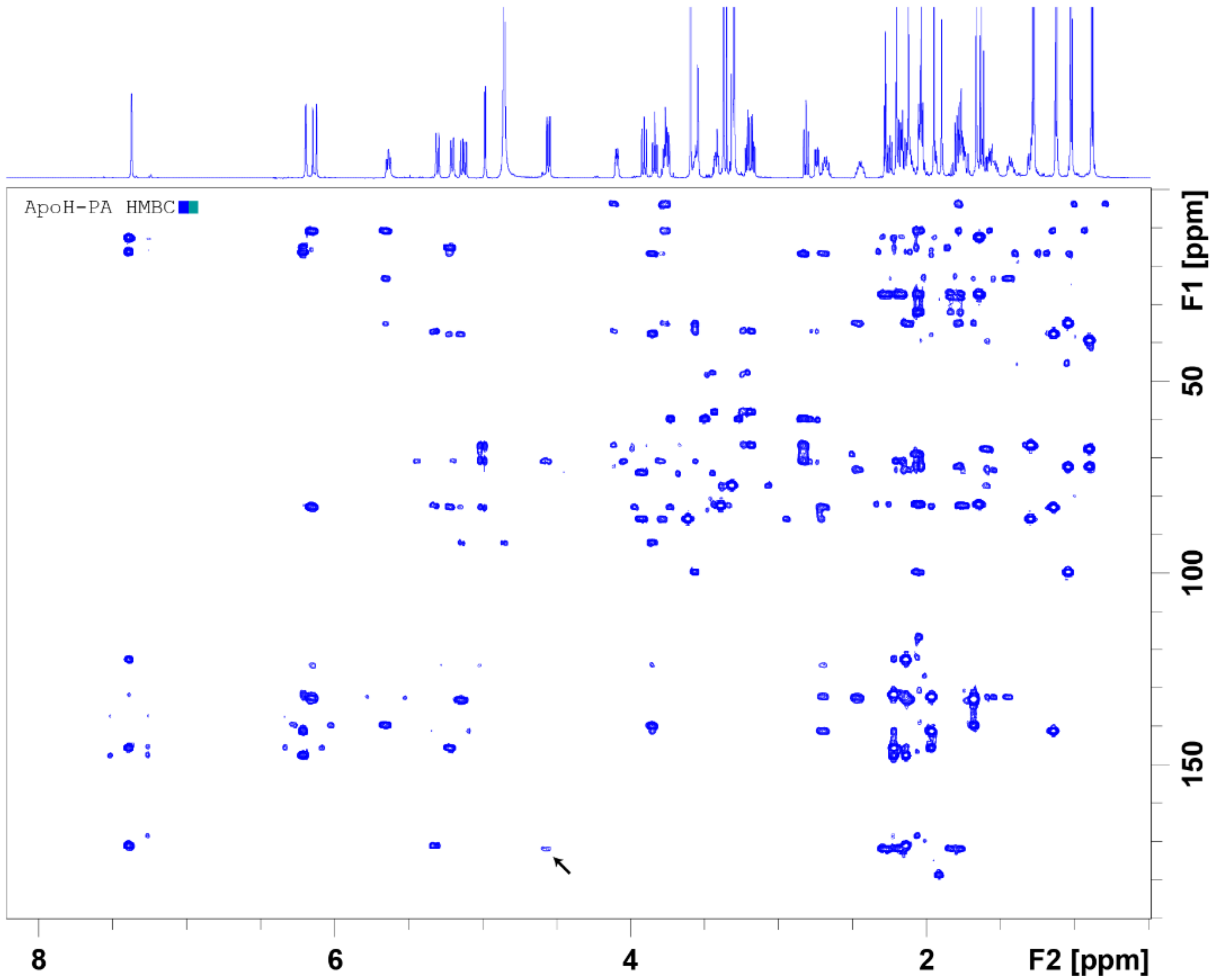

**Fig. S4 | HMBC of Apoptolidin H PA (2) in MeOD at 600 MHz**

HMBC spectrum of apoptolidin H PA in MeOD at 600 MHz with correlation between C2' and 1* (carbonyl of diazirine/alkyne ester) denoted by the arrow.

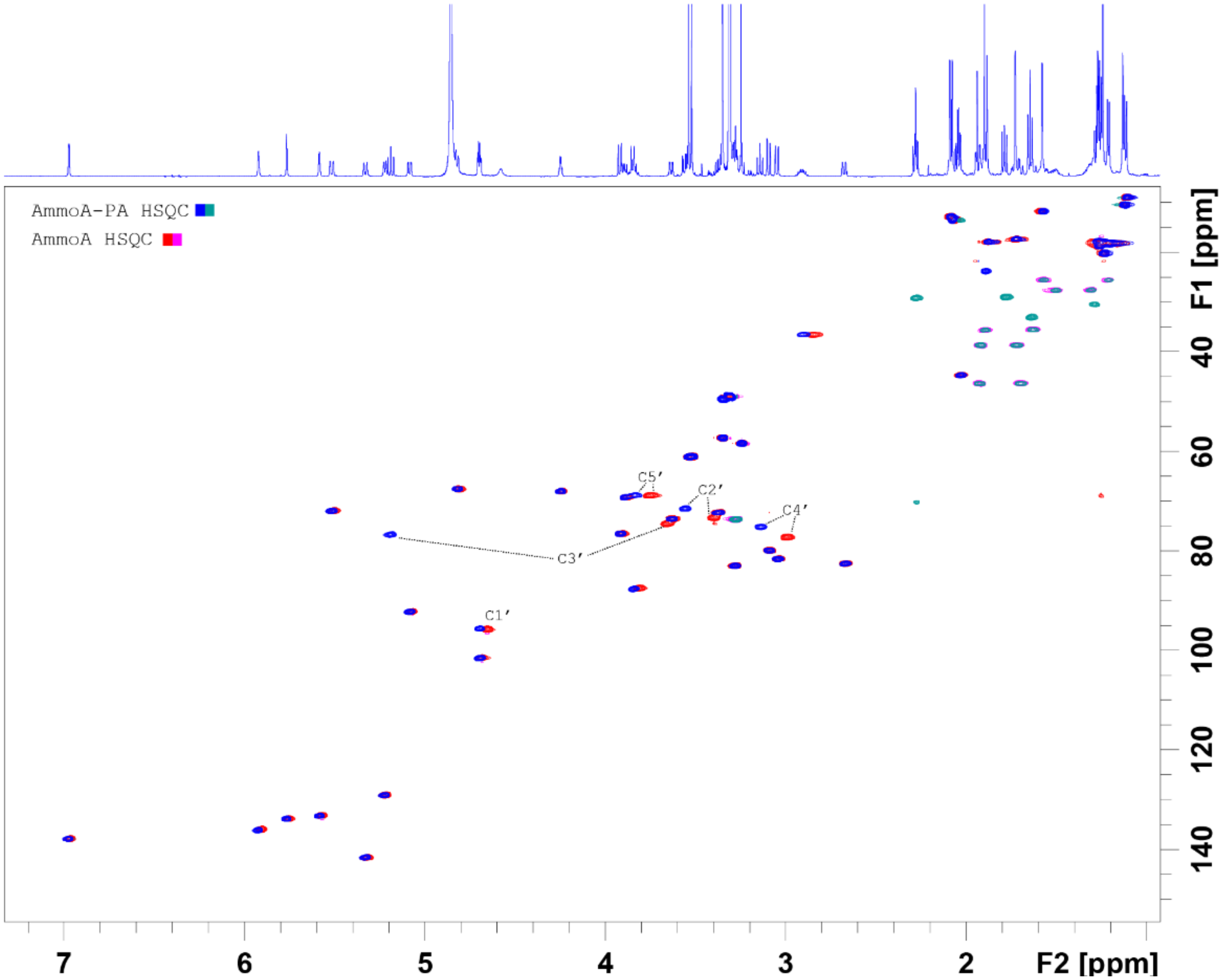

**Fig. S5 | HSQC of Ammocidin A PA (2) in MeOD at 600 MHz**

Ammocidin A PA (blue/green) on Ammocidin A (red/pink). C9 6-deoxy-α-L-glucose carbons labeled

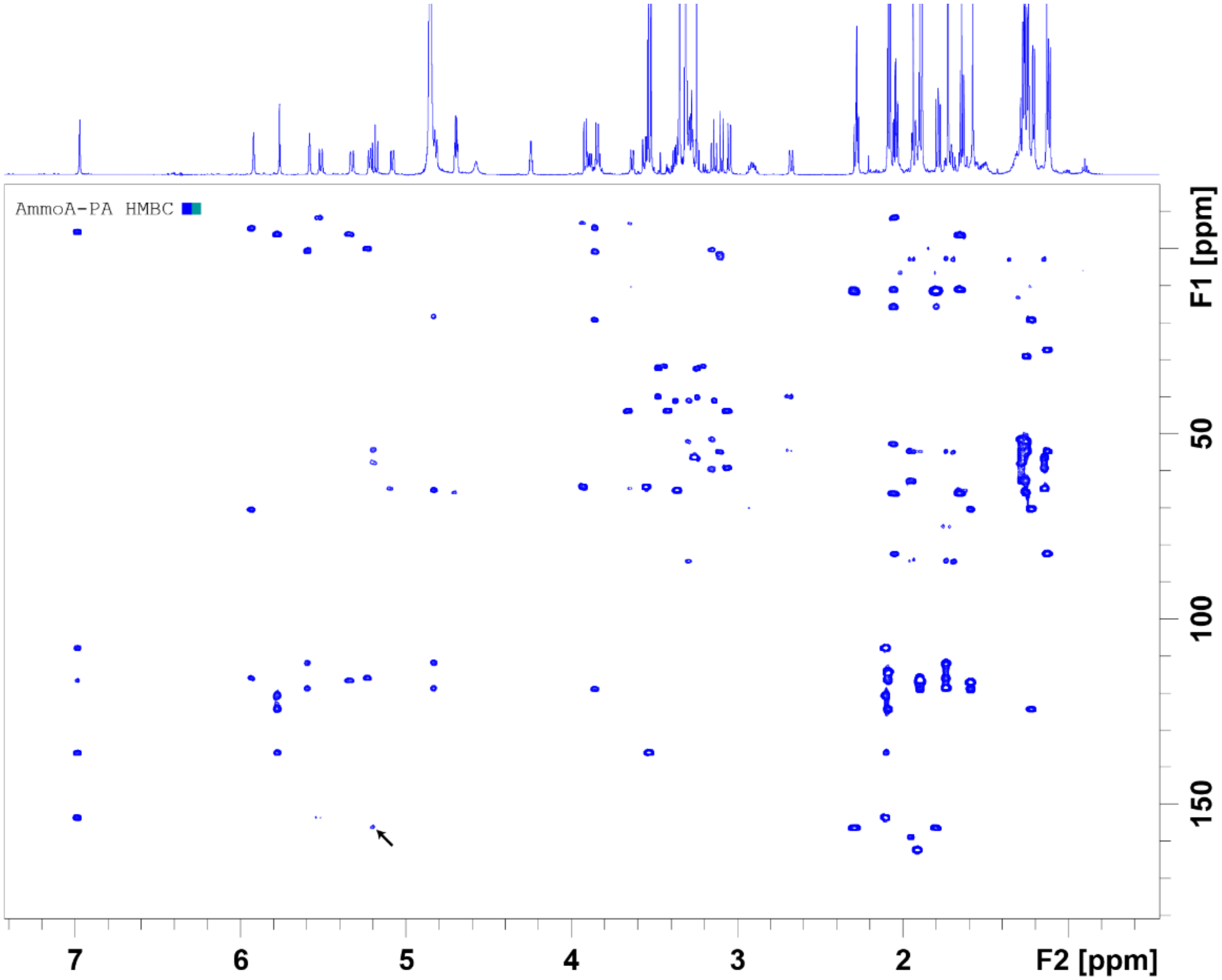

**Fig. S6 | HMBC of Ammocidin A PA (2) in MeOD at 600 MHz**

HMBC spectrum of ammocidin A PA in MeOD at 600 MHz with correlation between C3’ and 1* (carbonyl of diazirine/alkyne ester) denoted by the arrow.

**Supplementary Material References:**

1 Du, Y. *et al.* Biosynthesis of the Apoptolidins in Nocardiopsis sp. FU 40. *Tetrahedron* **67**, 6568-6575, doi:10.1016/j.tet.2011.05.106 (2011).

2 Baldwin, C. Biological and chemical properties of aurovertin, a metabolic product of Calcarisporium abuscula. *Lloydia* **27**, 88-95 (1964).

3 Krasnoff, S. B. & Gupta, S. Identification and directed biosynthesis of efrapeptins in the fungus *Tolypocladium geodes* gams (Deuteromycotina: Hyphomycetes). *J Chem Ecol* **17**, 1953-1962, doi:10.1007/BF00992580 (1991).

4 DeGuire, S. M. *et al.* Fluorescent probes of the apoptolidins and their utility in cellular localization studies. *Angew. Chem., Int. Ed.* **54**, 961-964, doi:10.1002/anie.201408906 (2015).
